# Hormonal regulation of the BRC1-dependent strigolactone transcriptome involved in shoot branching responses

**DOI:** 10.1101/2020.03.19.999581

**Authors:** Stephanie C. Kerr, Alexandre de Saint Germain, Indeewari M. Dissanayanke, Michael G. Mason, Elizabeth A. Dun, Christine A. Beveridge, Milos Tanurdzic

## Abstract

The plant hormone strigolactone (SL) inhibits shoot branching by suppressing the growth of axillary buds. This is thought to occur largely via regulation of the transcription factor BRANCHED1 (BRC1). Here, we clarify the central role of BRC1 and identify additional transcriptional responses by which SL might regulate axillary bud outgrowth in garden pea (*Pisum sativum*). We used a transcriptomic approach to identify differentially expressed transcripts in pea axillary buds in response to a synthetic SL, *rac*-GR24. Changes in transcript abundance were confirmed by measuring their response to GR24^5DS^. BRC1 was required for the regulation of over half of the fourteen GR24^5DS^-regulated genes, confirming its role as a mediator of SL transcriptional responses in axillary buds. All, but one, of the BRC1-dependent GR24^5DS^-regulated genes were also regulated by branch-promoting treatments cytokinin (CK) and/or decapitation in an opposing manner to SL. This suggests that SL, CK, and decapitation regulate shoot branching via a common pathway. We used correlational analyses of gene co-expression data to infer a gene regulatory network consisting of nine key co-expression modules correlated with *rac*-GR24 treatment. Enrichment of GO terms such as cell proliferation, carbohydrate responses, and abscisic acid and jasmonic acid hormone pathways suggest a role for these in SL-mediated inhibition of shoot branching. In summary, we have shown that BRC1 is indeed a key transcriptional regulator of the SL signalling pathway in pea buds as well as a focal point of the SL, CK and decapitation signalling pathways to coordinate shoot branching in pea buds.

**One Sentence Summary:** Identification of genes that are strigolactone-responsive and BRC1-dependent in pea buds reveals a high degree of overlap among strigolactone, cytokinin and decapitation response pathways.

## INTRODUCTION

Plants adapt their shoot architecture in response to changes in the environment, and this plasticity is regulated by particular hormones and signals which act together to repress or stimulate the growth of axillary buds to form branches. Among the plant hormones that regulate axillary bud growth, strigolactone (SL) and auxin inhibit it, while cytokinin (CK) and sugars promote it (Barbier et al., 2019b). The exact mechanisms by which each of these hormones and signals control the growth of axillary buds into branches and the ways in which their signalling pathways interact is still unclear. For SL responses, only a few studies have investigated the large-scale transcriptomic responses to SL (Mashiguchi et al., 2009; Mayzlish-Gati et al., 2010; Lantzouni et al., 2017; Zhan et al., 2018; Zha et al., 2019), and none have specifically examined these transcriptomic responses in axillary buds in intact plants. Determining how SL inhibits bud outgrowth will enable a better understanding of the molecular processes at play in deciding whether an axillary bud remains dormant or grows into a branch. This in turn will enable us to fine-tune axillary bud growth to create desirable shoot architectures in plants.

The signalling pathway of SLs from perception to repression of bud outgrowth has been only partially elucidated. The repressive effects of SL are triggered when it is perceived by an α/β hydrolase receptor (OsD14 (DWARF14); AtD14; PsRMS3 (RAMOSUS3)) along with an F-box protein (OsD3; AtMAX2 (MORE AXILLARY GROWTH2); PsRMS4), that is itself part of a SKP-CULLIN-F-BOX (SCF) complex (Waters et al., 2017). Upon its perception by the D14-MAX2-SCF complex, SL recruits one or more target proteins, including DWARF53 (D53) in rice and its homologs in Arabidopsis, the SUPPRESSOR OF MAX2-LIKE (SMXL) proteins, for ubiquitination and degradation by the 26S proteasome (Lopez-Obando et al., 2015). These D53/SMXL target proteins, in conjunction with the transcriptional co-repressors TOPLESS (TPL) and TOPLESS-RELATED (TPR), are likely to themselves repress downstream targets of the SL signalling pathway (Ma et al., 2017). Once SL-induced degradation of D53/SMXL proteins has taken place, these downstream transcriptional targets can be de-repressed, causing the inhibition of axillary bud outgrowth. Using mutant analyses, D53 and three members of the SMXL family in Arabidopsis—SMXL6, SMXL7 and SMXL8—have been implicated in the regulation of bud outgrowth. Dominant gain-of-function *smxl6, smxl7* or *d53* mutants that are unable to be ubiquitinated and degraded by SL have increased branching phenotypes (Jiang et al., 2013; Zhou et al., 2013; Wang et al., 2015; Liang et al., 2016), while loss-of-function triple *smxl6,7,8* mutants are able to completely rescue branching to wild-type (WT) levels in SL-mutant backgrounds (Soundappan et al., 2015; Wang et al., 2015). This implies that SL signalling for regulation of bud outgrowth likely occurs solely through D53/SMXL6,7,8. However, the precise number and nature of direct targets of the D53/SMXL proteins are still to be discovered.

Two main downstream targets of SL, and likely direct targets of D53/SMXL, have previously been characterised: the transcription factor BRC1, and the auxin transporter protein PIN1. BRC1 is a bud-specific TEOSINTE BRANCHED 1, CYCLOIDEA, PCF1 (TCP) transcription factor whose expression is correlated with inhibition of axillary bud outgrowth (reviewed in Barbier et al., 2019). *BRC1* is upregulated by SL treatment (Mashiguchi et al., 2009; Braun et al., 2012; Dun et al., 2012) and *brc1* mutants exhibit an increased branching phenotype that cannot be rescued by SL treatment (Brewer et al., 2009; Minakuchi et al., 2010; Braun et al., 2012). In wheat, D53 prevents SQUAMOSA PROMOTER-BINDING PROTEIN-LIKE (SPL) proteins from transcriptionally activating the *BRC1* ortholog, *TB1* (Liu et al., 2017). It is yet unknown whether SPLs in other species, including dicots, are involved in the regulation of *TB1/BRC1* by D53/SMXL. In Arabidopsis, BRC1 was recently shown to increase levels of abscisic acid (ABA) in buds by regulating genes involved in ABA biosynthesis and response, including the ABA biosynthesis gene 9-cis-epoxycarotenoid dioxygenase3 (NCED3) (González-Grandío et al., 2017). However, regulation of these genes by BRC1 could account for some, but not all, of the inhibitory effects of BRC1 on bud outgrowth. Therefore, other genes are likely to act downstream of BRC1 to regulate bud outgrowth.

PIN1 proteins are involved in the directional transport of auxin out of cells (Domagalska and Leyser, 2011). Polarisation of PIN1 proteins in axillary buds induces the formation of vascular strands from the bud to the polar auxin transport stream (PATS) in the main stem which is required for sustained axillary bud outgrowth (Balla et al., 2011; Chabikwa et al., 2019) that occurs after an initial growth period. The polarised auxin transport by PIN1 between the bud and PATS is aided by PIN3, 4 and 7 which facilitate movement of auxin through connective auxin transport (CAT) (Bennett et al., 2016). SL has been shown to decrease the localisation of PIN1 to the plasma membrane, which would reduce auxin transport and inhibit bud growth (Shinohara et al., 2013). Supporting a role of PIN1 acting downstream of SL, branching in SL mutants is correlated with elevated *PIN1* expression, increased PIN1 localisation to the plasma membrane, and increased auxin transport (Bennett et al., 2006).

Though BRC1 and PIN1 are important targets of SL, their actions do not entirely explain the inhibition of branching by SL. *BRC1* expression is not always correlated with bud inhibition (Arite et al., 2007; Seale et al., 2017), and preventing auxin transport out of buds using an auxin transport inhibitor, 1-N-Naphthylphthalamic acid (NPA), does not inhibit early bud growth (Brewer et al., 2009; Brewer et al., 2015; Chabikwa et al., 2019). This indicates the possibility that other targets of SL are yet to be discovered.

SL regulates many developmental processes in different tissues throughout the plant, and downstream SL signalling differs between these processes. For example, SL decreases PIN1 polarisation in shoots, but increases PIN2 polarisation in roots (Koltai, 2014). Previous studies identifying downstream transcript or protein targets of SL used whole seedlings (Mashiguchi et al., 2009; Chen et al., 2014; Li et al., 2014; Lantzouni et al., 2017), roots (Mayzlish-Gati et al., 2010), shoots (Zhan et al., 2018), or buds of decapitated plants (Zha et al., 2019), but it is likely that SL targets differ in other tissues and conditions. This has likely contributed to some SL-regulated genes remaining unidentified.

Antagonistic to SL response in buds, the plant hormone CK is a promoter of branching and is also thought to act partially via BRC1 (Braun et al., 2012; Dun et al., 2012). Although the exact mechanisms by which CK achieves this also remains unclear, it may involve cell cycle regulation as CK promotes cell division in other tissues such as the shoot apical meristem (SAM) (Müller and Leyser, 2011). The mobile sugar sucrose is another important long-distance signal that promotes axillary bud outgrowth in plants (Barbier et al., 2015b). Recent work has demonstrated that the initial decapitation-induced bud outgrowth induced by removal of rapidly expanding leaves is likely caused by sucrose (Mason et al., 2014; Barbier et al., 2015a), with auxin and other hormones contributing to the regulation of continued bud growth (Barbier et al., 2019b). Interestingly, *BRC1* has also been shown to be transcriptionally regulated by both decapitation and sucrose treatments (Mason et al., 2014; Gao et al., 2016; Moreno-Pachon et al., 2017), suggesting that BRC1 may act as an integrator of multiple branching signals.

In order to identify downstream targets of SL specifically related to the regulation of bud outgrowth, we profiled transcriptomic changes in pea axillary buds in response to a single SL treatment known to cause a rapid and long-term inhibition of bud outgrowth. Pea was selected as the model species as it possesses large axillary buds separated by long internodes enabling easy treatment and harvest of bud-specific tissue. A racemic mixture of synthetic SL, *rac*-GR24 (or (+/-)-GR24), composed of GR24^5DS^ (or (+)-GR24), which perfectly mimics the configuration of natural SL, and GR24^*ent*-5DS^ (or (-)-GR24), was supplied to buds which were then collected over a six hr time course and subjected to RNA sequencing (Kerr et al., 2017). We inferred SL-responsive gene networks using weighted gene co-expression network analysis (WGCNA) and identified genes whose SL regulation is dependent on BRC1. To assess whether BRC1 acts as an integrator of multiple branching signals (such as CK and shoot tip removal) to regulate downstream transcriptional targets, we examined whether our *BRC1*-dependent SL-regulated genes were also responsive to CK and decapitation treatments in pea axillary buds.

## RESULTS

### *rac*-GR24 transcriptionally regulates few genes in axillary buds prior to outgrowth

To define the time frame for early SL-induced changes in bud outgrowth, we performed timeseries measurements on *rms5* SL-deficient plants, using node 2 buds treated with 1 μM of the synthetic SL, *rac*-GR24. *rms5* mutant plants were used as they produce insufficient SL to inhibit bud outgrowth, enabling the use of exogenous *rac*-GR24 application to inhibit active node 2 bud growth. further

The first measurable effect of exogenous *rac*-GR24 on *rms5* node 2 bud growth was observed at 6.5 hrs post-treatment (Figure S1A). We observed upregulation of *PsBRC1* expression as early as 1 hr after treatment with *rac*-GR24 to node 2 buds of *rms5* plants using qRT-PCR (Figure S1B). Consequently, in order to capture the gene expression changes occurring over this timeframe of the early stages of bud growth inhibition by SL, a 4-point time course between 1 hr and 6 hrs was utilised for RNA sequencing (Kerr et al., 2017). Pairwise comparisons were made between the mock and *rac*-GR24 treatments at each time point following treatment, and a generalised linear model (GLM) was used to identify transcripts showing statistically significant changes in steady-state transcript levels between the mock- and *rac-GR24* treated samples across the 6 hr time course.

Using a false discovery rate (FDR) threshold of 0.05 we discovered only a small number of statistically significantly differentially expressed (DE) transcripts, 14 in total (Table S1 and Supplementary Dataset 1) at any given time point. This included no DE transcripts at 1 and 2 hrs after treatment, even though we demonstrated that *rac-*GR24 can regulate *PsBRC1* expression as early as 1 hr using qRT-PCR (Figure S1B). To determine the accuracy of the DE analyses, qRT-PCR was performed on all 14 transcripts and a validation rate of 79% was observed (Table S1). One reason for the relative paucity of DE genes is likely the result of biological variation in individual pea buds and their responses to *rac-*GR24. To identify additional prospective *rac-*GR24-regulated transcripts we relaxed the FDR threshold to 0.5, which identified a further 292 putative *rac-*GR24-regulated transcripts (Table S1 and Supplementary Dataset 1). qRT-PCR analyses performed on 48 DE transcripts validated half of them as expected by the FDR threshold (Table S1).

In total, 26 transcripts representing 24 genes were validated as *rac-*GR24-regulated by qRT-PCR (Table S1); two genes, *PsERF061* and *PsSMXL8* were represented by two partial transcripts, comp114246_c0 and comp57631_c0, and comp86930_c0 and comp86930_c1, respectively, as annotated in Kerr et al. (2017). To address the lack of DE genes identified at early time points in the RNA-seq analysis, despite evidence that *PsBRC1* can be regulated by *rac*-GR24 at 1 hr, we tested the 24 validated *rac-*GR24-regulated genes using qRT-PCR at all four time points following *rac*-GR24 treatment. We were able to demonstrate that many of these genes were significantly differentially expressed at time points not identified as statistically significantly by the RNA-seq analyses, including three genes at 1 hr and seven genes at 2 hrs (Figure 1).

**Figure 1.**
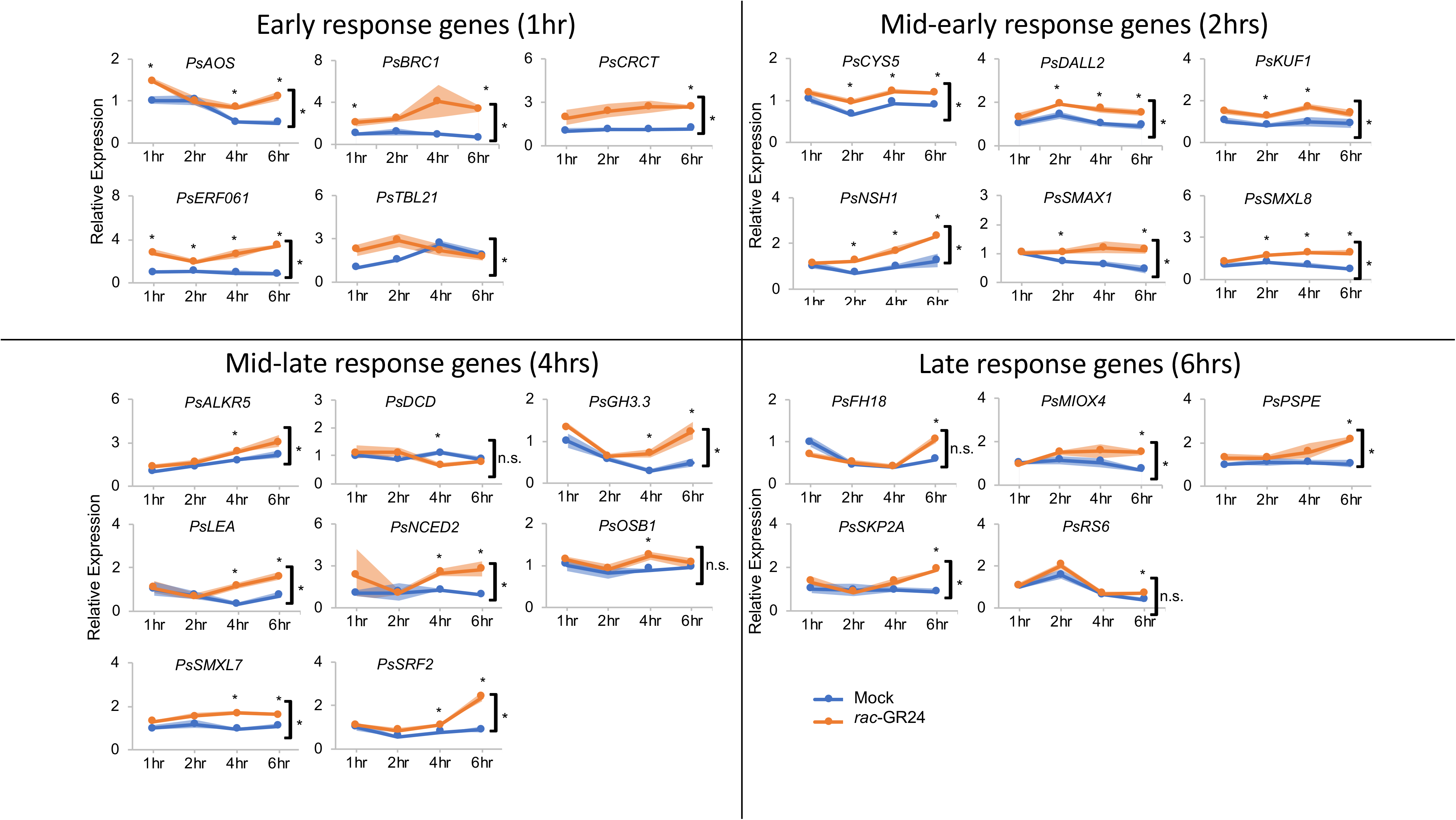
Expression of SL-regulated genes 1, 2, 4 or 6 hrs after *rac*-GR24 treatment. The bud at node 2 of 7-day-old *rms5-3 Pisum sativum* plants was treated for 1, 2, 4 or 6 hrs with a solution containing 0 (Blue) or 1 μM (Orange) *rac*-GR24. Expression determined using qRT-PCR is represented relative to the mock treatment at 1 hr and was normalized against the geomean of three internal reference genes: *EF1α*, *GADPH* and *TUB2*. Data are means ± SE (n = 3-4 pools of ~30-40 buds). * denotes means significantly different from the mock treatment for each time point (Student t-test; P<0.05) or over all time points (denoted by the bracket; two-way ANOVA; P<0.05; n.s. denotes means not significantly different from the mock treatment). The genes were divided into four groups (Early, Mid-early, Mid-late and Late) based on the earliest time at which they responded to *rac*-GR24 treatment (1hr, 2hrs, 4hrs and 6hrs, respectively).

Of the 24 confirmed *rac-*GR24-regulated genes, only two were previously annotated in pea in GenBank, comp78442_c0 *(PsBRC1;* GenBank: AEL12230.1; Braun et al., 2012) and comp70495_c0 *(PsNCED2; 9-cis-epoxycarotenoid dioxygenase 2;* GenBank: BAC10550.1), while the other 22 were annotated by sequence similarity to an Arabidopsis protein (Table 1; Kerr et al., 2017). This process identified genes already known to be SL responsive in other species, such as *PsBRC1* (comp78442_c0; Braun et al., 2012) and two pea orthologs of *SUPPRESSOR OF MAX2-LIKE* genes (Stanga et al., 2013); *PsSMXL7* (comp78374_c1) and *PsSMXL8* (comp86930_c0). Additionally, the closest homolog of *PsNCED2* (comp70495_c0) in Arabidopsis, *AtNCED3* (Rodrigo et al., 2006), has previously been shown to be SL-responsive (González-Grandío et al., 2017).

**Table 1.** SL-regulated gene annotation. Annotations of the 24 SL-regulated pea transcripts by sequence comparison using BLAST or by closest ortholog assignment using OrthoDB (Kriventseva et al., 2015). Gene descriptions/names bolded are those used to describe each pea transcript.

| Pea transcript | Best BLAST annotation from the pea genome (from Kreplak et al., 2019) | Best BLAST annotation from the nr protein database (from Kerr et al., 2017) |  | Best BLAST annotation from *Arabidopsis thaliana* (from Kerr et al., 2017) |  | Closest ortholog (OrthoDB) |  | Designated pea name |
| --- | --- | --- | --- | --- | --- | --- | --- | --- |
|  |  | Gene description/name | Locus tag [Species] | Gene description/name | Locus tag | Gene description | Locus tag [Species] |  |
| comp102807_c0 | Psat7g207240 | fatty acid hydroperoxide lyase; allene oxide synthase | MTR_4g068550 [*Medicago truncatula*] | ALLENE OXIDE SYNTHASE (AOS); CYTOCHROME P450 74A (CYP74A); DELAYED DEHISCENCE 2 (DDE2) | AT5G42650 | fatty acid hydroperoxide lyase; allene oxide synthase | MTR_4g068550 [*Medicago truncatula*] | PsAOS (Allene Oxide Synthase) |
| comp114246_c0 | Psat0s2105g0240 | AP2 domain class transcription factor | MTR_5g009410 [*Medicago truncatula*] | integrase-type DNA-binding superfamily protein; ETHYLENE RESPONSIVE TRANSCRIPTION FACTOR 61 (ERF061) | AT1G64380 | no orthologs | no orthologs | PsERF061 (Ethylene Responsive transcription Factor 61) |
| comp378909_c0 | Psat2g158920 | no BLAST hit | n/a | no BLAST hit | n/a | late embryogenesis abundant (LEA) domain protein | MTR_5g020520 [*Medicago truncatula*] | PsLEA (Late Embryogenesis Abundant domain protein) |
| comp54718_c1 | Psat4g039000 | probable galactinol--sucrose galactosyltransferase 6 isoform X2 | LOC101497789 [*Cicer arietinum*] | RAFFINOSE SYNTHASE 6 (RS6); DARK INDUCIBLE 10 (DIN10); | AT5G20250 | no orthologs | no orthologs | PsRS6 (Raffinose Synthase 6) |
| comp55397_c0 | Psat4g009240 | actin-binding FH2 (formin 2) family protein | MTR_4g131020 [*Medicago truncatula*] | Actin-binding FH2 (Formin Homology) protein; Formin-like protein 18 (FH18) | AT2G25050 | actin-binding FH2 (formin 2) family protein | MTR_4g131020 [*Medicago truncatula*] | PsFH18 (Formin-like protein 18) |
| comp55781_c0 | Psat7g113960 | plant-specific domain<br>TIGR01615<br>family protein | MTR_4g088770<br>[*Medicago truncatula*] | sugar phosphate exchanger,<br>putative (DUF506) | AT4G32480 | uncharacterised<br>proteins | GLYMA07G31690<br>/<br>GLYMA13G24760<br>[*Glycine max*] | PsPSPE<br>(Putative Sugar<br>Phosphate<br>Exchanger) |
| comp56263_c0 | Psat7g172080 | phospholipase<br>A1-Igama3,<br>chloroplastic | LOC101496884<br>[*Cicer arietinum*] | alpha/beta-Hydrolases<br>superfamily protein;<br>phospholipase A1-Igama3;<br>DAD1-Like Lipase 2<br>(DALL2) | AT1G51440 | phospholipase<br>A1 | MTR_4g077070<br>[*Medicago truncatula*] | PsDALL2<br>(DAD1-Like<br>Lipase 2) |
| comp70495_c0 | Psat1g001480 | nine-cis-<br>epoxycarotenoi<br>d dioxygenase<br>2 (PsNCED2) | n/a<br>[*Pisum sativum*] | NINE-CIS-EPOXYCAROTENOID<br>DIOXYGENASE 3<br>(NCED3); SALT<br>TOLERANT 1 (STO1);<br>SUGAR INSENSITIVE 7<br>(SIS7) | AT3G14440 | no orthologs | no orthologs | PsNCED2<br>(Nine-Cis-Epoxycarotenoi<br>d Dioxygenase<br>2) |
| comp70728_c0 | Psat3g034440 | inosine-uridine<br>preferring<br>nucleoside<br>hydrolase | MTR_7g104270<br>[*Medicago truncatula*] | NUCLEOSIDE<br>HYDROLASE 1 (NSH1);<br>URIDINE-RIBOHYDROLASE 1<br>(URH1) | AT2G36310 | no orthologs | no orthologs | PsNSH1<br>(Nucleoside<br>Hydrolase 1) |
| comp71047_c1 | Psat5g077440 | uncharacterize<br>d protein | LOC101507417<br>[*Cicer arietinum*] | CCT motif family protein | AT5G53420 | CCT motif<br>protein | MTR_3g091340<br>[*Medicago truncatula*] | PsCRCT (CO2-Responsive<br>CONSTANS,<br>CONSTANS-like, and Time<br>of Chlorophyll<br>a/b Binding<br>Protein) |
| comp73484_c0 | Psat1g197600 | Pmr5/Cas1p<br>GDSL/SGNH-like acyl-<br>esterase family<br>protein | MTR_2g016060<br>[*Medicago truncatula*] | TRICHOME<br>BIREFRINGENCE-LIKE 21<br>(TBL21) | AT5G15890 | uncharacterised<br>proteins | GLYMA13G07200<br>/<br>GLYMA19G05770<br>/<br>GLYMA18G51491 | PsTBL21<br>(Trichome<br>Birefringerence-Like 21) |
|  |  |  |  |  |  |  | [*Glycine max*] |  |
| comp74214_c2 | Psat6g011160 | organellar single-stranded DNA-binding protein | MTR_1g021080<br>[*Medicago truncatula*] | ORGANELLAR SINGLE-STRANDED DNA-BINDING PROTEIN 1 (OSB1) | AT1G47720 | - | - | PsOSB1 (Organellar Single-Stranded DNA-Binding protein 1) |
| comp74856_c2 | Psat2g101440 | probable indole-3-acetic acid-amido synthetase GH3.1 | LOC101507657<br>[*Cicer arietinum*] | Indole-3-acetic acid-amido synthetase GH3.3 | AT2G23170 | GH3 protein | GLYMA02G13910<br>[*Glycine max*] | PsGH3.3 (Indole-3-acetic acid-amido synthetase) |
| comp78374_c1 | Psat5g158160 | double Clp-N motif P-loop nucleoside triphosphate hydrolase superfamily protein | MTR_3g464150<br>[*Medicago truncatula*] | double Clp-N motif-containing P-loop nucleoside triphosphate hydrolases superfamily protein; SUPPRESSOR OF MAX2-LIKE 7 (SMXL7) | AT2G29970 | uncharacterised proteins | GLYMA11G27120 /<br>GLYMA18G06990<br>[*Glycine max*] | PsSMXL7 (SUPPRESSOR OF MAX2-LIKE 7) |
| comp78442_c0 | Psat4g043560 | **PsBRC1**; TCP family transcription factor | n/a<br>[*Pisum sativum*] | TCP family transcription factor 18 (TCP18); BRANCHED1 (BRC1) | AT3G18550 | PsBRC1; TCP family transcription factor | n/a<br>[*Pisum sativum*] | PsBRC1 (BRANCHED1) |
| comp78689_c0 | Psat4g010720 | uncharacterised protein | LOC101508071<br>[*Cicer arietinum*] | double Clp-N motif-containing P-loop nucleoside triphosphate hydrolases superfamily protein; SUPPRESSOR OF MAX2 1 (SMAX1) | AT5G57710 | no orthologs | no orthologs | PsSMAX1 (SUPPRESSOR OF MAX2-1) |
| comp80781_c0 | Psat2g143160 | F-box/kelch-repeat protein SKIP25, putative | MTR_5g026580<br>[*Medicago truncatula*] | F-box/kelch-repeat protein SKIP25; KAR-UP F-box 1 (KUF1) | AT1G31350 | F-box/kelch-repeat protein SKIP25, putative | MTR_5g026580<br>[*Medicago truncatula*] | PsKUF1 (KAR-UP F-box 1) |
| comp81043_c1 | Psat5g198080 | protein STRUBBELIG-RECEPTOR FAMILY 2-like | LOC101498175<br>[*Cicer arietinum*] | no BLAST hit | n/a | STRUBBELIG-RECEPTOR FAMILY 2 (SRF2) | AT5G06820<br>[*Arabidopsis thaliana*] | PsSRF2 (Strubbelig-Receptor Family 2) |
| comp81735_c1 | Psat1g013720 | DCD (development and cell death) domain protein | MTR_2g066940<br>[*Medicago truncatula*] | DCD (Development and Cell Death) domain protein | AT2G35140 | no orthologs | no orthologs | PsDCD (Development and Cell Death domain protein) |
| comp86930_c0 | Psat5g129160 | uncharacterised protein | LOC101500860<br>[*Cicer arietinum*] | double Clp-N motif-containing P-loop nucleoside triphosphate hydrolases superfamily protein; SUPPRESSOR OF MAX2-LIKE 8 (SMXL8) | AT2G40130 | uncharacterised proteins | GLYMA11G35410<br>[*Glycine max*] | PsSMXL8 (SUPPRESSOR OF MAX2-LIKE 8) |
| comp87019_c0 | Psat4g004160 | cysteine proteinase inhibitor | MTR_4g133540<br>[*Medicago truncatula*] | cystatin-like protein; CYSTEINE PROTEINASE INHIBITOR (CYS5) | AT5G47550 | cysteine proteinase inhibitor | MTR_4g133540<br>[*Medicago truncatula*] | PsCYS5 (Cysteine proteinase inhibitor 5) |
| comp87415_c1 | Psat6g039800 | F-box SKP2-like protein | MTR_1g029500<br>[*Medicago truncatula*] | F-box/RNI-like superfamily protein; SKP2-like protein 1 (SKP2A) | AT1G21410 | - | - | PsSKP2A (SKP2-like Protein 1) |
| comp90222_c0 | Psat3g188960 | aldo/keto reductase family oxidoreductase | MTR_7g021680<br>[*Medicago truncatula*] | NAD(P)-linked oxidoreductase superfamily protein; Probable aldo-keto reductase 5 (ALKR5) | AT1G60730 | aldo/keto reductase family oxidoreductase | MTR_7g021820/<br>MTR_7g021670<br>[*Medicago truncatula*] | PsALKR5 (Probable Aldo-Keto Reductase 5) |
| comp93400_c0 | Psat7g019040 | myo-inositol oxygenase | MTR_8g102150<br>[*Medicago truncatula*] | MYO-INOSITOL OXYGENASE 4 (MIOX4) | AT4G26260 | myo-inositol oxygenase | MTR_8g102150<br>[*Medicago truncatula*] | PsMIOX4 (Myo-Inositol Oxygenase 4) |

### GR24^5DS^ and *rac*-GR24 treatment to pea buds elicit similar gene expression responses

*rac*-GR24 is a racemic mixture of two enantiomers, GR24^5DS^ and GR24^ent-5DS^, harbouring the same stereochemistry to the endogenous SL, 5-deoxystrigol (5DS), and its non-natural stereoisomer, *ent-5DS*, respectively (Scaffidi et al., 2014). These GR24 enantiomers can have different effects on plant growth and development with GR24^5DS^ eliciting similar responses to endogenous SLs, likely because GR24^5DS^ preferentially signals through the SL receptor, D14, while GR24^*ent*-5DS^ preferentially signals through the karrikin receptor, KARRIKIN INSENSITIVE 2 (KAI2) (Scaffidi et al., 2014; Li et al., 2016). We confirmed that GR24^5DS^, similar to *rac*-GR24, was able to significantly reduce node 2 bud growth in *rms1* seedlings over 1 week following treatment, while karrikin, KAR_1_, was unable to reduce bud growth as seen previously for SL biosynthesis mutants in Arabidopsis (Nelson et al., 2011; Figure S2).

We tested responses of the 24 *rac*-GR24-regulated genes to GR24^5DS^ treatment after 6 hrs in *rms1* buds, and found that at least 14 of these genes showed a more than 1.5-fold response to GR24^5DS^ (Figure 2), although it should be noted that only 20 of the 24 genes were regulated by *rac*-GR24 at 6 hrs (Figure 1). This correlation between *rac*-GR24 and GR24^5DS^ regulation of gene expression was confirmed by clustering analysis of gene expression responses measured by qRT-PCR, where 6 hr treatment with GR24^5DS^ clustered closely with 4 and 6 hr treatment with *rac*-GR24 (Figure 3). The six genes that did not respond to GR24^5DS^ treatment at 6 hrs, despite previously responding to *rac*-GR24 treatment at 6 hrs, were *PsFH18, PsRS6, PsGH3.3, PsCYS5, PsMIOX4*, and *PsLEA*. The lack of GR24^5DS^ response by these six genes could be due to experimental variation, but nevertheless suggests that SL regulation of these transcripts is unlikely to be important for the regulation of bud growth as GR24^5DS^ could inhibit bud growth in the absence of responses by these five genes, at least measured at 6 hrs following treatment. It is possible that these six genes could be regulated by the other component of *rac*-GR24, GR24^*ent*-5DS^, which was not tested here. However, we did test the effect of KAR_1_, a karrikin molecule that signals through the same receptor as GR24^*ent*-5DS^, KAI2 (Scaffidi et al., 2014). Most of the *rac*-GR24-regulated genes were not affected by KAR_1_ treatment, as expected for SL-specific response genes (Figure 2). In fact, clustering analysis of the 24 *rac*-GR24-regulated genes demonstrated that KAR_1_ treatment clustered more closely with CK and decapitation treatment than with SL treatment (Figure 3).

**Figure 2.**
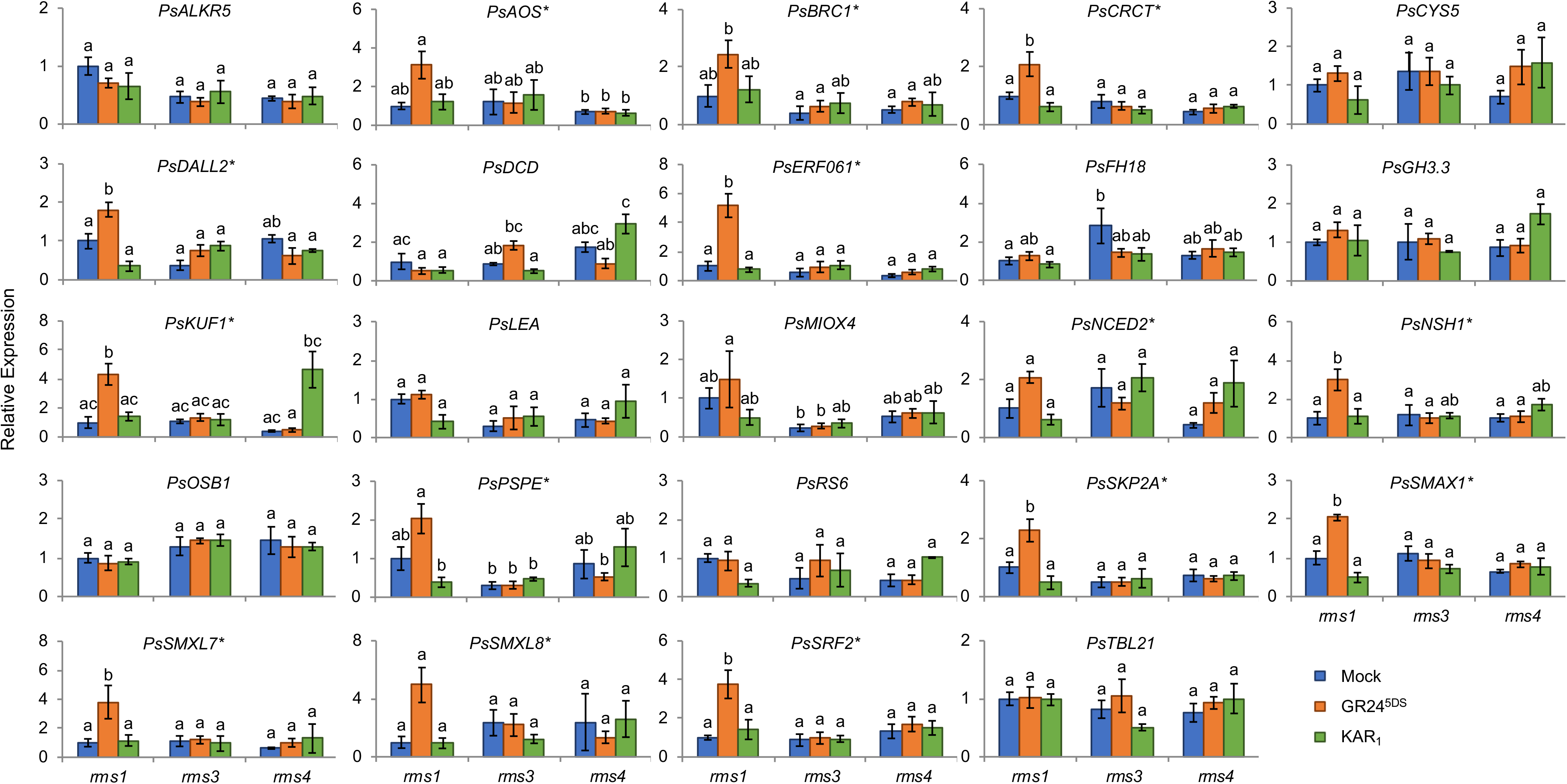
GR24^5DS^ regulation of SL-regulated genes is RMS3- and RMS4-dependent. The bud at node 2 of 9-day-old *rms1-2, rms3-2* and *rms4-1 Pisum sativum* plants was treated for 6 hrs with 0 (Blue) or 1 μM (Orange) GR24^5DS^ or 1 μM (Green) kAR_1_. Expression is represented relative to the *rms1-2* mock treatment and was normalized against the geomean of three internal reference genes: *EF1α*, *GADPH* and *TUB2*. Data are means ±SE (n = 3 pools of ~10-20 buds). Data were analysed using a two-way ANOVA with Tukey comparisons of means; different letters represent statistical differences of P<0.05. * indicates genes that were classified as GR24^5DS^-regulated for further analysis in Figures 4, S5 and S6.

**Figure 3.**
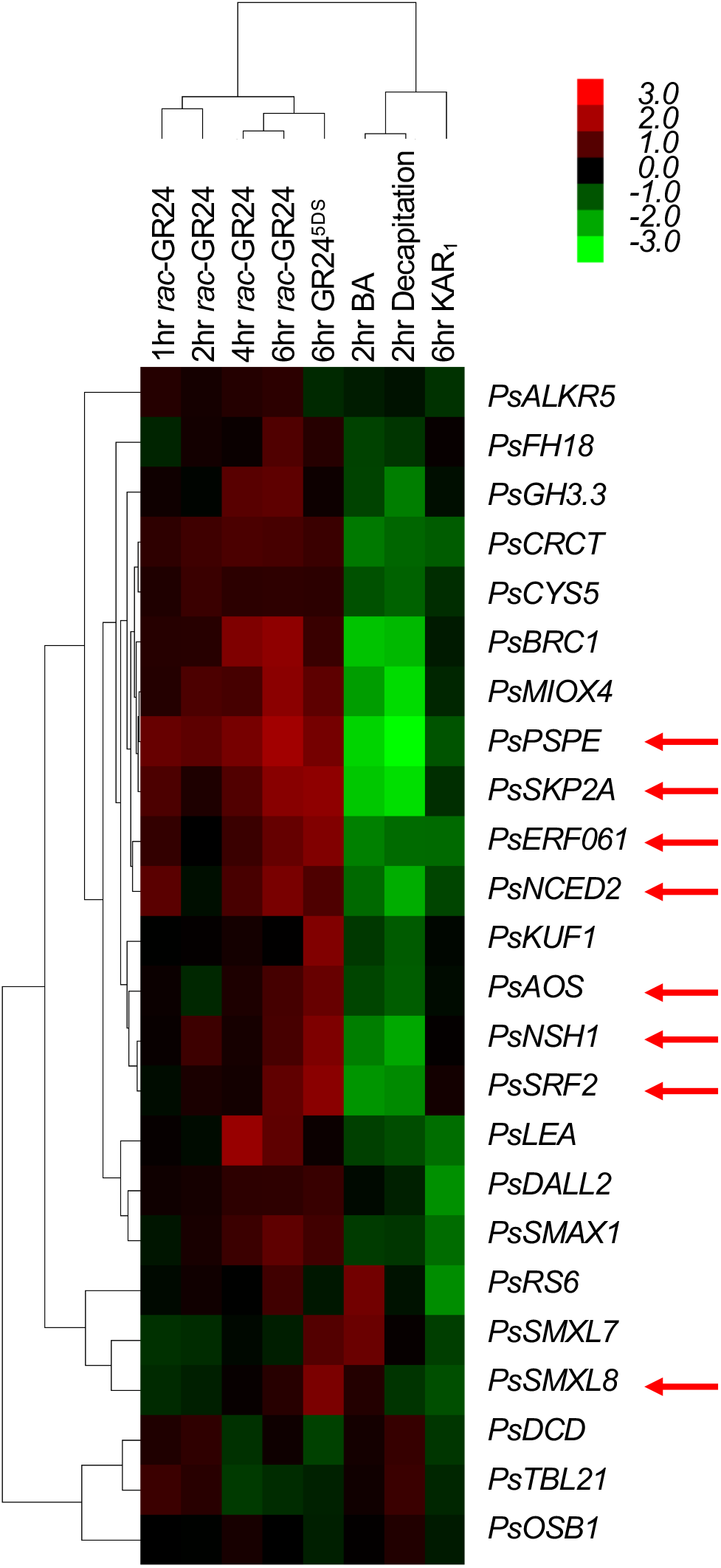
Heatmap representing the difference in fold change of transcript levels in mock versus treated node 2 pea buds. *rac-*GR24 data is from Figure 1, GR24^5DS^ and KAR1 data is from Figure 2, BA data is from Figure S5, and Decapitation data is from Figure S6. Red arrows indicate genes that were also found to be regulated by *rac*-GR24 in a BRC1-dependent manner as determined in Figure 5.

To determine if any of the fourteen GR24^5DS^-regulated genes we identified play a major role in shoot branching, we examined branching phenotypes in Arabidopsis T-DNA insertion mutants of most of the GR24^5DS^-regulated genes. We choose to examine mutant phenotypes in Arabidopsis due to the easy accessibility of mutant lines in Arabidopsis compared to pea. Branching phenotypes of mutants of *BRC1* (Hubbard et al., 2002; Aguilar-Martínez et al., 2007; Finlayson, 2007; Minakuchi et al., 2010; Braun et al., 2012), *NCED3* (Reddy et al., 2013; González-Grandío et al., 2017), *CRCT*(Morita et al., 2015) and *SMXL7, SMXL8* and *SMAX1* (Jiang et al., 2013; Zhou et al., 2013; Soundappan et al., 2015; Wang et al., 2015) have previously been characterised in other species, and so were not further analysed here.

Most of the T-DNA mutants we examined did not display a measurable branching phenotype (Figure S3), perhaps because they only play a minor, or redundant, role in shoot branching and therefore do not display a significant branching phenotype on their own. For example, in Arabidopsis AtSMXL6, AtSMXL7 and AtSMXL8 act somewhat redundantly in the positive regulation of shoot branching downstream of SL perception. Subsequently, only double or triple *Atsmxl6, Atsmxl7* and/or *Atsmxl8* mutants can significantly alter the branching phenotype in SL mutant backgrounds (Soundappan et al., 2015; Wang et al., 2015). Additionally, single, double and triple *Atsmxl6, Atsmxl7* and/or *Atsmxl8* mutants have no effect on branching on a WT background (Soundappan et al., 2015; Wang et al., 2015) as WT plants already produce so few branches that increasing SL signalling by removing AtSMXL6, AtSMXL7 and/or AtSMXL8 cannot further reduce branching. In our analysis, only one mutant, *Atskp2a-1*, displayed an observable branching phenotype in our growth conditions (Figure S3). Further experiments suggest that AtSKP2A may negatively regulate branching (Supplementary Dataset 2), but additional investigation is required to provide conclusive results.

### Gene co-expression analysis reveals key processes and regulatory factors responsive to SL

We took advantage of the variation in gene expression response to *rac*-GR24 over the time series by employing weighted gene co-expression network analysis (WGCNA; Langfelder and Horvath, 2008) to infer a gene co-expression network and identify gene co-expression modules significantly correlated with *rac*-GR24 treatment. To do this, we first filtered genes whose raw expression values across all samples were equal to or higher than 10 (from Kerr et al., 2017), and then selected the top 10% (5000) genes sorted by their variance-stabilised expression values in ascending order across 23 RNA samples. Twenty gene co-expression modules were identified among these top 5000 variable genes using *rac*-GR24 treatment and treatment length as covariables (Figure S4). Of these, nine modules showed significant correlation (P<0.05) with the *rac*-GR24 treatment and/or treatment length, comprising 1891 of the 5000 genes. These nine modules (4, 6, 9, 10, 12, 15, 17, 18, and 19) showed a high degree of correlated co-expression between, as well as, within modules (Figure 4A).

**Figure 4.**
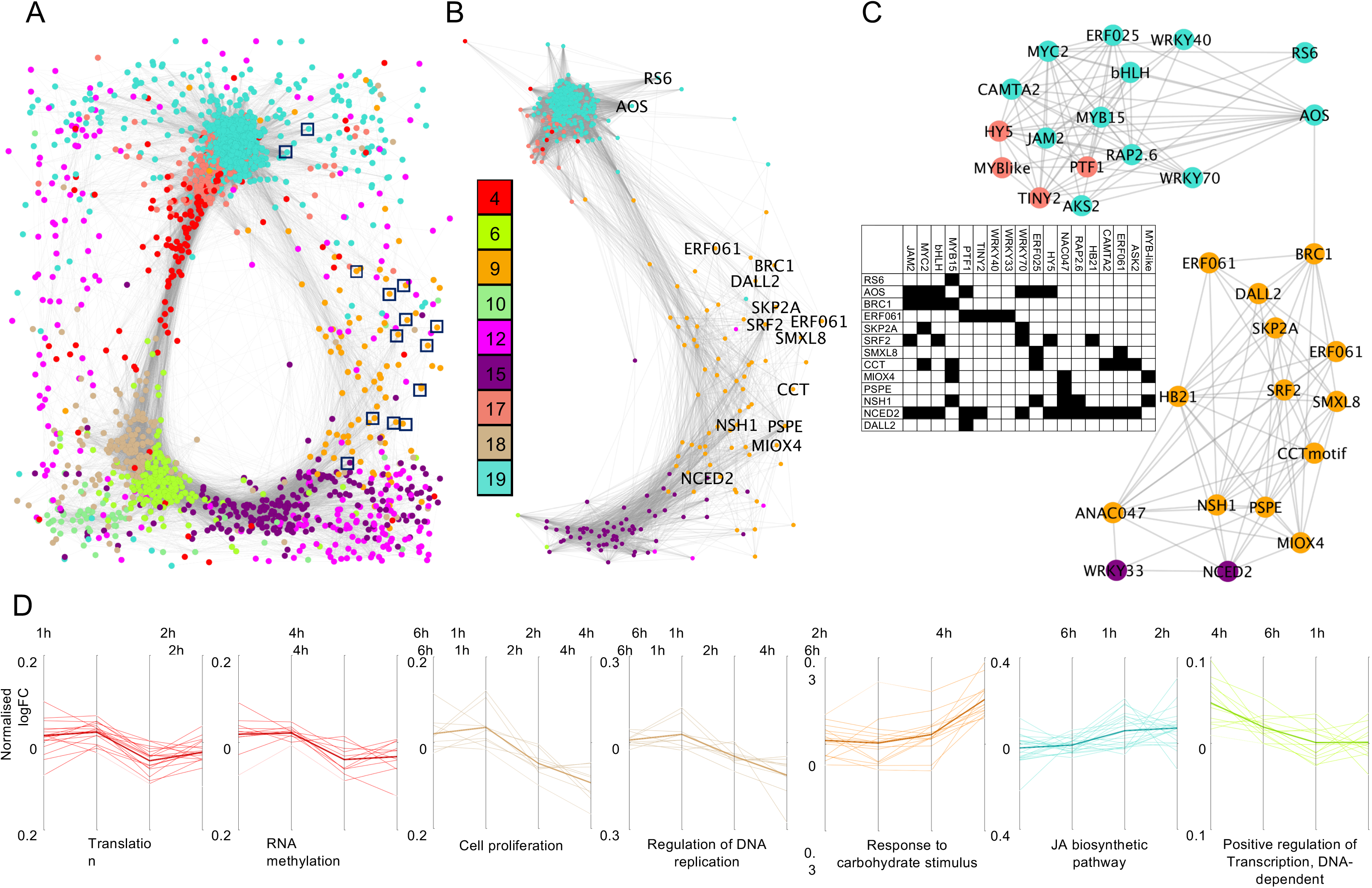
*rac*-GR24-regulated gene co-expression network. **A)** Cytoscape visualization of the nine gene coexpression modules (listed in Table S3) showing significant correlation with 1 μM *rac*-GR24 treatment; genes are shown as nodes and the co-expression correlation between nodes as grey lines. The squares highlight the 13 *rac*-GR24-regulated genes from the text (also shown in part B). B) Subset of co-expression network consisting of the 13 *rac*-GR24-regulated genes present in the network and their first neighbors (direct co-expression correlation edges). Edges shown in red represent connections with transcription factors within the sub-network nodes (further highlighted in C). C) The transcription factor first neighbors of the 13 SL-regulated genes (identities shown) and the predicted TF-target relationships between them (inset table) using TF2Network predictions (Kulkarni et al., 2018). D) Time-course profile of normalized gene expression profiles of module-specific genes with a particular GO term enrichment (listed underneath each profile). Gene expression profile for all genes within the GO term stated under the figure; Y axis represents the variance-stabilized normalized log fold change (logFC) values between *rac*-GR24 and Control treatment at each time point; line colors match the color of the module they were derived from; the darker line represents the average expression values of all genes in that dataset.

We examined the enriched Biological Process GO terms present in these modules to explore the functional relevance of the gene co-expression correlations across the nine modules (Kerr et al., 2017; Supplementary Dataset 3). Many of the enriched GO terms within each module are functionally related. For example, module 4 was enriched in translation, ribosome biogenesis, and RNA methylation processes (Supplementary Dataset 3 and Figure 4D), while module 17 was enriched in circadian rhythm and light-related GO terms (Supplementary Dataset 3). These results point to coordinated regulation of several processes involved in growth and development by SL, including downregulation of genes involved in translation, RNA methylation, cell proliferation and regulation of DNA replication, and overall up-regulation of genes involved in response to carbohydrate stimulus, jasmonic acid (JA) biosynthesis and positive regulation of transcription (Figure 4D). The high level of connectivity between gene co-expression modules 9 and 19 (Figure 4A, 4B) and their functional classifications (Supplementary Dataset 3) in combination with the temporal dynamics of expression (Figure 4D) infer that hormone biosynthetic and response pathways are regulated in the same timeframe as responses to carbohydrate stimulus.

To better understand the roles of the 24 confirmed *rac*-GR24-regulated genes we identified if any were present as nodes within the co-expression network. Of the 24 *rac*-GR24-regulated genes, 13 were present within three of the nine significant modules: PsRS6, PsAOS, PsBRC1, PsERF061, PsSKP2A, PsSRF2, PsSMXL8, PsCRCT, PsMIOX4, PsPSPE, PsNSH1, PsNCED2, and PsDALL2 (Figure 4A, 4B), mostly within module 9 (orange in Figure 4). Furthermore, these 13 *rac*-GR24 genes were connected to 434 first neighbours, or 23% of the genes present in the nine modules, and comprised 29% of the edges present in the network (Figure 4B), suggesting that a significant portion of genes located within the nine co-expression modules are coexpressed with confirmed SL-regulated genes.

To test if any of the co-expression correlation between the 13 *rac*-GR24 genes and their 434 direct nodes in the pea co-expression network may represent a conserved genetic interaction in other species, we compared it with publicly available data from Arabidopsis using gene coexpression, protein-DNA, and protein-protein interaction data in GeneMANIA (Warde-Farley et al., 2010). We were able to recover ~10-20% of the pea nodes within the Arabidopsis networks for seven of the 13 pea genes (Table S2). This result highlights conserved interaction candidates for SL-regulated genes across flowering plant lineages.

To identify SL-regulated transcription factors (TFs), in addition to BRC1, we investigated the TFs present in the network. Of the 1191 TFs that were identified in the pea bud transcriptome (Kerr et al., 2017), 92 were present in the nine significant modules (data not shown). As might be expected from a gene co-expression network, a high proportion of nodes (60%) and edges (85%) from the nine significant modules were associated with the 92 TFs, suggesting that these TFs may be involved in the regulation of most of the genes present in the nine significant modules. In support of this, we identified several TFs as the top 5 hub genes in three of the modules: PsNAC047 and PsHB21 in module 9, PsDREB3 and PsHY5 in module 17, and PsWRKY18 and PsERF113 in module 18 (Supplementary Dataset 4). These six TFs were only a subset of the 49 TFs also present within the sub-network of genes co-expressed with the 13 *rac*-GR24-regulated genes (the 434 node sub-network, Figure 4B). The 49 TFs were co-expressed with 419 (97%) genes in this sub-network (Figure 4B). Several TFs within this group have previously been implicated in SL signalling, most importantly the TF PsHB21 (González-Grandío et al., 2017). We found PsHB21 co-expressed with 68 genes, including PsBRC1 (Figure 4B, 4C). While

BRC1 has been shown to regulate HB21 in Arabidopsis (González-Grandío et al., 2017), our data are not able to distinguish between direct and indirect interactions between PsBRC1 and PsHB21, but raise a strong possibility that this component of the SL signalling pathway is conserved between Arabidopsis and pea.

Publicly available Arabidopsis TF-target gene promotor binding predictions (Kulkarni et al., 2018) provided further information on the likely directionality of some of the network edges. The Arabidopsis homologues of 18 TFs from the sub-network had their binding sites significantly enriched within the 1kb of promoter sequence of the 434 gene sub-network (Figure 4C). These observations provide initial information on the possible hierarchy of transcriptionally regulated steps in SL signalling. For example, the transcription factor HB21 has been previously identified as a direct target of BRC1 while itself regulating ABA responses through AtNCED3 (González-Grandío et al., 2017). Here we provide indirect evidence of a similar pathway in pea, with PsHB21 showing co-expression correlation with PsBRC1 and PsNCED2 (likely ortholog of AtNCED3). Importantly, PsHB21 also shows a likely role in the regulation of, or by, the JA signalling pathway through its interactions with PsJAM2 and bHLH (Figure 4C). The TF binding motifs of the JA signalling TFs JAM2, MYC2, and bHLH, on the other hand, have been identified (most often all three together) in the promoters of 6 of the 13 *rac*-GR24 genes, including BRC1 19/03/2020 10:20:00 PM. These observations further strengthen the temporal order of SL responses implied by gene expression profiles and co-expression correlations between the genes in modules 9 and 19 (Figure 4).

### All GR24^5DS^-responsive genes require the α-β hydrolase RMS3 and the F-box protein RMS4 for their SL response

The effect of GR24^5DS^ treatment on gene expression was examined in *rms3* and *rms4* buds. RMS3 and RMS4 are both involved in perception of SL (Gomez-Roldan et al., 2008; de Saint Germain et al., 2016). As expected, all fourteen genes that were responsive to GR24^5DS^ in the SL deficient *rms1* mutant were no longer responsive to GR24^5DS^ in either the *rms3* or *rms4* mutants (Figure 2), confirming that these genes require the canonical RMS3 and RMS4 components of the SL perception and signalling pathway.

### PsBRC1 is required for regulation of some, but not all, SL-regulated genes

To elucidate whether PsBRC1 is required for SL regulation of gene expression, we examined expression of the fourteen GR24^5DS^-regulated genes after *rac*-GR24 treatment in node 2 buds of the loss-of-function *Psbrc1* mutant (Braun et al., 2012). As *rac*-GR24 has minimal effect on gene expression in wild type (WT) plants, likely due to high endogenous SL levels, we conducted this experiment on an *rms1* background, comparing gene expression responses in *rms1* buds to those of *brc1 rms1* buds.

As expected, SL responses were observed for all fourteen GR24^5DS^-regulated genes in the *rms1* mutant at 6 hrs after treatment (Figure 5). Over half of the genes were not responsive to SL when BRC1 was mutated in the *rms1* mutant (Figure 5), including *PsAOS, PsERF061, PsNCED2, PsNSH1, PsPSPE, PsSKP2A, PsSRF2* and *PsSMXL8*, and were designated as BRC1-dependent SL-regulated genes. It should be noted that although *rac*-GR24 was unable to significantly increase *PsSMXL8* expression in the *rms1 brc1* double mutant, there was still a more than 1.5-fold increase in *PsSMXL8* expression (Figure 5), which may suggest that *PsSMXL8* is only partially dependent on BRC1 for *rac*-GR24 responsiveness. The remaining six genes were still responsive to SL in the *brc1* mutant (Figure 5), including *PsBRC1, PsCRCT, PsDALL2, PsKUF1, PsSMAX1* and *PsSMXL7*, and were therefore designated as BRC1-independent SL-regulated genes. Such a BRC1-independence could theoretically include genes that are required for SL-regulated BRC1 response or are completely BRC1 independent.

**Figure 5.**
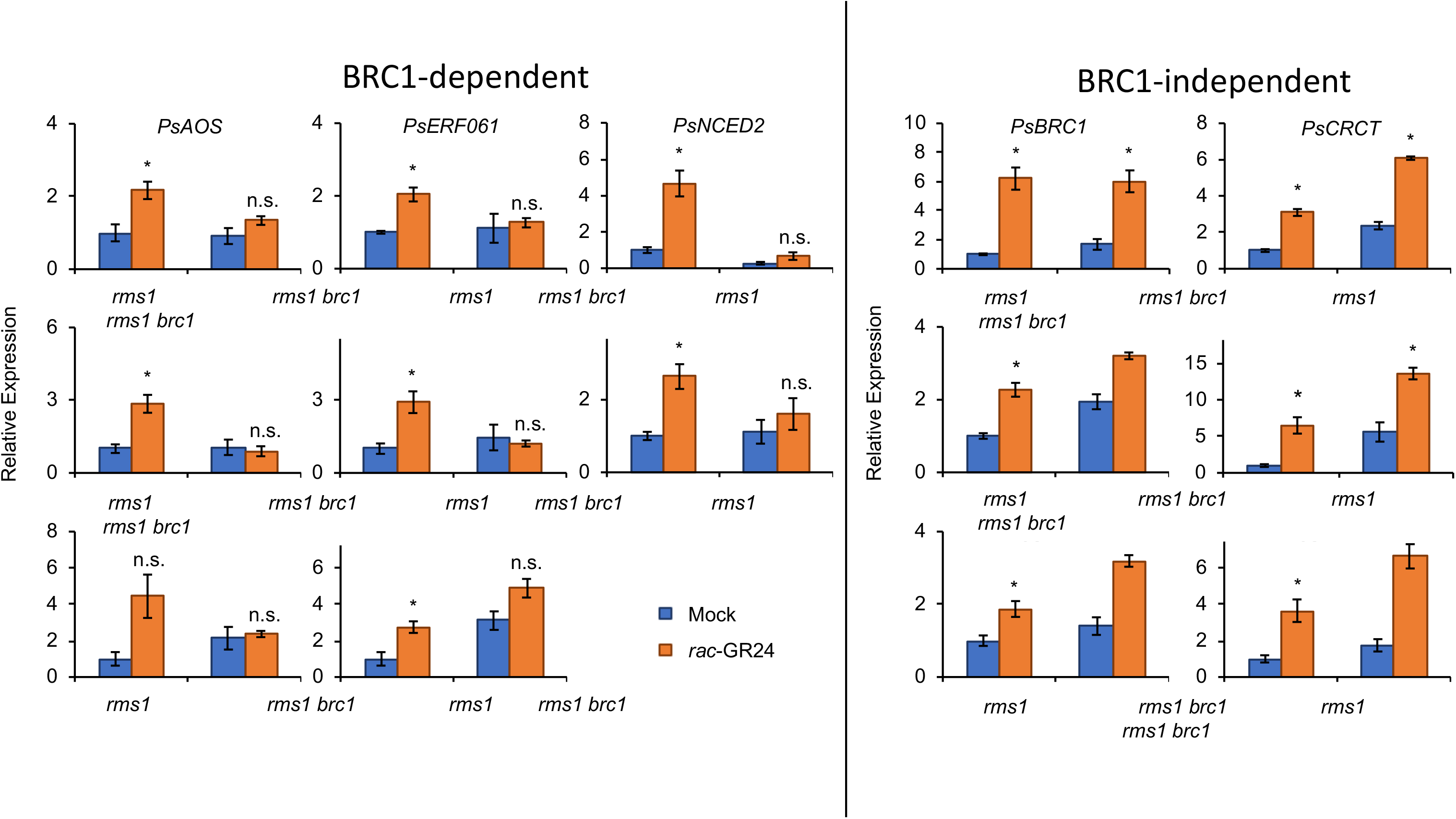
PsBRC1 is required for SL regulation of some SL-regulated genes. The bud at node 3 on 13-day-old *rms1-10* and *rms1-10 PsBRC1^Té^ Pisum sativum* seedlings was treated for 6 hrs with containing 0 (Blue) or 1 μM (Orange) *rac*-GR24. Expression determined using qRT-PCR is represented relative to the mock treatment in BRC1 and was normalized against the geomean of three internal reference genes: *EF1α*, *GADPH* and *TUB2*. Data are means ± SE (n = 3 pools of ~30-40 buds). * denotes means significantly different from the mock treatment for each genotype (Student t-test; P<0.05; n.s. denotes means not significantly different from the mock treatment).

### CK and decapitation also regulate many SL-regulated genes

BRC1 is also regulated by other long-distance signals, including, but not limited to, CK and sucrose (Braun et al., 2012; Dun et al., 2012; Mason et al., 2014). We therefore used the synthetic CK, 6-benzyl aminopurine (BA) and decapitation of the shoot tip to test if these different pathways that modulate BRC1 expression also regulate the fourteen GR24^5DS^-upregulated genes. Nine of the fourteen GR24^5DS^-upregulated genes were downregulated by both CK and decapitation (Figures S5 and S6). This implies that a significant portion of the SL response pathway is antagonistically regulated by CK and decapitation, as demonstrated in Figure 3. All but one of the genes regulated in this manner were BRC1-dependent genes implying that this antagonistic regulation of gene expression by these regulators of shoot branching occurs through BRC1. Interestingly, given their role in SL signalling, *PsSMXL7* and *PsSMXL8* were the only two genes upregulated by both SL and CK (Figure S5), while *PsSMXL7* was the only gene upregulated by both SL and decapitation (Figure S6).

To see if CK and decapitation regulation of SL-responsive transcripts is also present in our SL-regulated gene co-expression network, we compared the list of 434 genes co-expressed with the 13 confirmed *rac*-GR24-regulated genes (Figure 4B) with published datasets of CK and decapitation-regulated genes in Arabidopsis. Of the 434 pea genes present in Figure 4B, 298 were annotated with an Arabidopsis gene (Kerr et al. 2017). For CK we found that 61 of the 298 genes in our network (20%; Hypergeometric test; P<4.8E-15) were also regulated by BA treatment after 2 hrs in Arabidopsis whole seedlings (Brenner et al., 2005), while for decapitation we found that 62 of the 298 genes in our network (21%; Hypergeometric test; P<3e-08) were also regulated in Arabidopsis axillary buds and leaves 24 hrs after decapitation of the main stem (Tatematsu et al., 2005). Not unexpectedly, a high proportion of the genes in the same module as the 13 confirmed *rac*-GR24-regulated genes (Module 9) were present in both the CK (25%) and decapitation (36%) datasets. Although these datasets are not exact replicas of the conditions we used for CK and decapitation regulation of genes in Figures S5 and S6 (e.g. CK was performed on whole seedlings rather than buds, while decapitation treatment was 24 hrs not 2 hrs), the fact that ~20% of genes that are co-expressed with confirmed *rac*-GR24-regulated genes in pea are also regulated by CK and decapitation in Arabidopsis suggests that CK and decapitation regulation of SL-responsive genes may be conserved across species.

## DISCUSSION

Regulation of shoot architecture is a key process that allows a plant to adapt to environmental conditions. This regulation involves multiple hormones and signals, including SL, that act in concert to determine whether axillary buds grow into branches and ultimately determine the shoot architecture of a plant. Here we have identified key transcriptional targets of SL in pea buds and have demonstrated that a core set of genes respond to multiple signals through the transcription factor BRC1. By examining the functions of these core set of genes we can further our understanding of how SL inhibits axillary bud outgrowth.

### Identification of novel BRC1-dependent and -independent SL-response genes in pea buds

Not unexpectedly, three of the five genes classified as early *rac*-GR24 response genes are transcription factors *(PsBRC1, PsERF061* and *PsCRCT;* Figure 1). In addition, the GO term ‘positive regulation of transcription’ was upregulated by *rac*-GR24 at 1 hr in Module 6 (Figure 4D). This indicates a temporal pattern of gene regulation where early-response transcription factors regulate the expression of late-response genes. We were able to test this hypothesis for PsBRC1 by examining transcriptional responses to *rac*-GR24 in *brc1* mutant buds. In fact, we found that over half (8) of the SL-regulated genes required BRC1 for a SL response (Figure 5). This confirms BRC1 as an essential component of downstream SL signalling and has informed a model of SL-regulation of genes in pea buds that highlights BRC1-dependent and BRC1-independent pathways (Figure 6). The identity of these BRC1-dependent genes by sequence similarity to other species includes genes likely involved in ABA biosynthesis (NCED3; Iuchi et al., 2001), JA biosynthesis (AOS; Laudert et al., 1996), cell cycle regulation (SKP2A; del Pozo et al., 2002), and nucleoside metabolism (NSH1; Jung et al., 2011). We suggest that these BRC1-dependent SL-regulated genes are good candidates for being involved in the regulation of shoot branching and require further investigation. Interestingly, the SL-regulated BRC1-independent genes included SMAX1, which has previously been suggested to be more specific to karrikin KAI2-mediated signalling than SL signalling, although a limited role for SMAX1 in shoot branching regulation was suggested (Soundappan et al., 2015).

**Figure 6.**
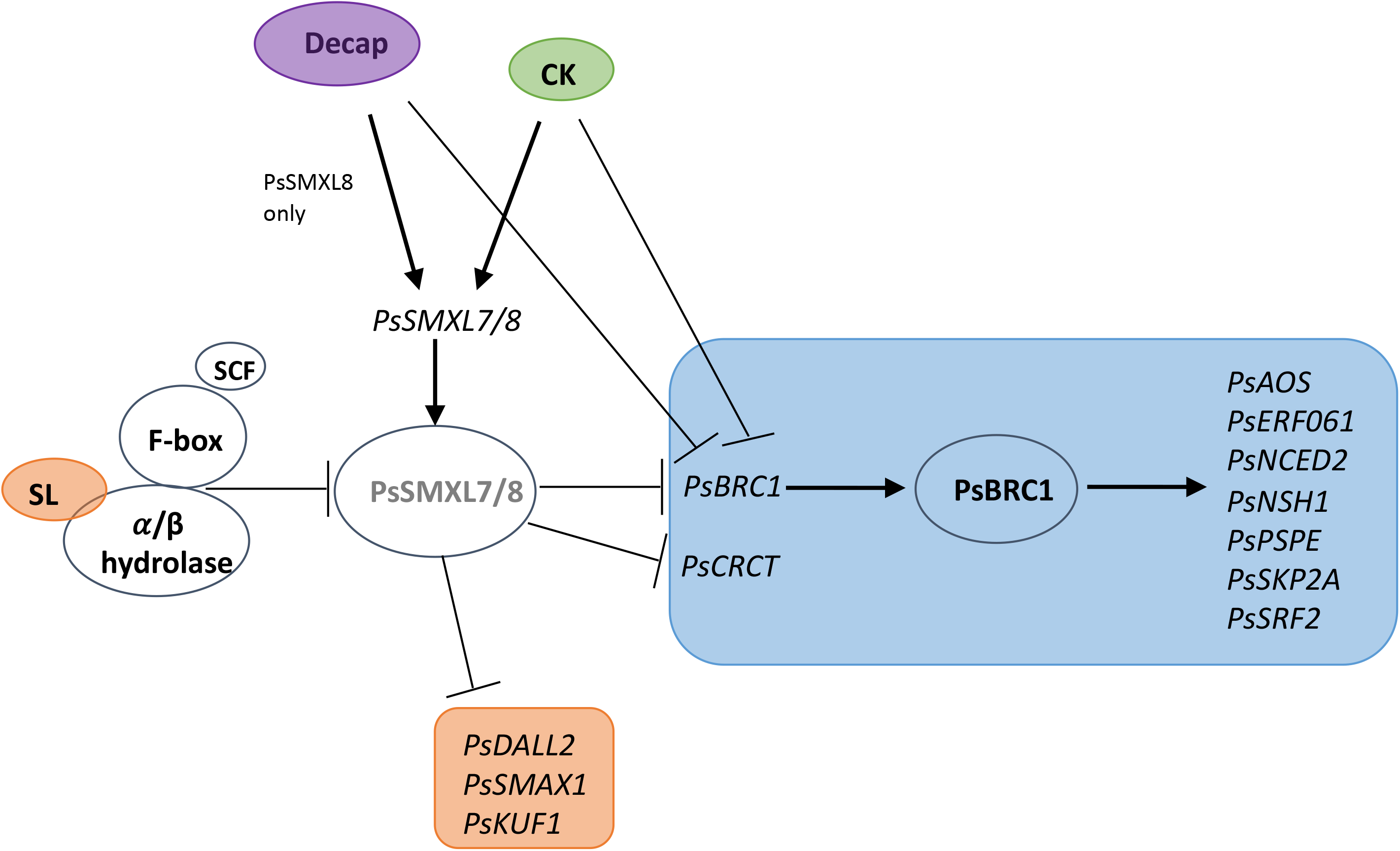
Summary of BRC1-dependent and BRC1-independent gene regulation by SL, CK and decapitation in pea buds. Genes in orange box indicate those regulated by SL, but not decapitation or cytokinin treatment, while genes in blue box indicate those regulated by SL, CK and decapitation.

ABA has long been associated with bud dormancy (e.g. Tucker, 1977), and more recently as a downstream target of BRC1 to inhibit axillary bud outgrowth (González-Grandío et al., 2017). Here we identified the putative ABA biosynthesis gene *PsNCED2* as a BRC1-dependent SL-regulated gene. The closest homologs of *PsNCED2* in Arabidopsis, *AtNCED3*, and maize, viviparous14 (vp14), are direct transcriptional targets of AtBRC1 and its homolog in maize, TB1 (González-Grandío et al., 2017; Dong et al., 2019). The *Atnced3* mutant exhibits an increased branching phenotype, especially in response to low R:FR light treatment (Reddy et al., 2013; Yao and Finlayson, 2015; González-Grandío et al., 2017), supporting a role for AtNCED3 in the regulation of shoot branching. AtBRC1 can directly regulate the HD-Zip genes, *AtHB21, AtHB40* and *AtHB53*, which themselves could also regulate *AtNCED3* expression (González-Grandío et al., 2017). It remains to be seen if BRC1 can directly regulate HD-Zip genes in other species such as pea, however, we identified two HD-Zip genes, annotated as HB13 and HB21 (Kerr et al., 2017), in our list of putative *rac*-GR24-regulated genes (Supplementary Dataset 1).

There is some evidence that JA negatively regulates shoot branching (Zavala and Baldwin, 2006; Zhong et al., 2013; Rubio-Moraga et al., 2014), although contrasting results (e.g. Liu and Finlayson, 2019) require further investigation of the role of JA in shoot branching. Soybean SL biosynthesis and signalling mutants have been shown to have reduced JA levels in leaves and altered expression of JA biosynthetic genes (Haq et al., 2017) supporting a potential interaction between JA and SL. SL regulation of JA is further supported by our GO term analysis (Supplementary Dataset 3) with enrichment of JA biosynthesis and signalling GO terms after *rac*-GR24 treatment in Modules 6, 15 and 19. JA and JA biosynthetic genes were also recently identified as targets of TB1 in rice (Dong et al., 2019). We identified PsAOS, which in Arabidopsis is a key enzyme involved in JA biosynthesis (Chehab et al., 2011), as a BRC1-dependent SL-regulated target gene. Interestingly, we identified another gene likely involved in JA biosynthesis, DAD1-Like Lipase 2 (DALL2) (Lin et al., 2016), that was upregulated by SL, but was BRC1-independent. Although *Ataos* or *Atdall2* mutants did not exhibit a branching phenotype under our growth conditions, a dominant mutation in *DONGLE*, a gene in the same AtPLA1-I family as *DALL2* has an increased branching phenotype in Arabidopsis (Hyun et al., 2008).

Some members of the TCP family of transcription factors to which BRC1 belongs control cell proliferation by regulating expression of cell-cycle progression genes (Nicolas and Cubas, 2016). The maize ortholog of BRC1, TB1, may establish bud dormancy by regulating cell proliferation genes (Dong et al., 2019). We have further identified a gene co-expression module responsive to *rac*-GR24 (Module 18) that is enriched in GO terms associated with the cell cycle (Figure 4D and Supplementary Dataset 3) that could provide potential BRC1-targets. Furthermore, homologs of two genes, PsSRF2 and PsSKP2A, that we confirmed as BRC1-dependent in their SL-response have putative roles in the cell cycle in other species; SKP2A is involved in the proteolysis of cell cycle transcription factors via the Ubiquitin/26S pathway in Arabidopsis (del Pozo et al., 2006), while SRF2 belongs to the same LRR-V family as STRUBBELIG (SUB), a gene involved in determining the orientation of the cell division plane as well as cell proliferation and morphogenesis, although whether SRF2 has a similar function is unknown (Chevalier et al., 2005; Eyüboglu et al., 2007).

### SL treatments in intact plants regulate a relatively small number of transcripts

Only a few studies investigating transcriptional responses to SL have been performed to date. Both Mashiguchi et al. (2009) and Lantzouni et al. (2017) investigated the early transcriptional responses to *rac*-GR24 in whole Arabidopsis seedlings and found only 64 and 73 SL-regulated transcripts, respectively. Of these genes six were in common with our list of 24 *rac*-GR24-regulated genes including KUF1, SMAX1, SMXL7, SMXL8, ERF061 and CRCT, which is significantly higher than expected by chance (Hypergeometric test; P<1.8e-11 and 4.96e-7, respectively). This suggests that SL regulation of these genes may not be specific to the bud, as their regulation by SL could be observed in whole seedlings. Another study by Zha et al. (2019) investigated transcriptional responses two hrs after *rac*-GR24 treatment in axillary buds in decapitated rice plants. They identified 1529 *rac*-GR24-regulated transcripts, most of which were upregulated. Only one transcript was in common with our list of confirmed *rac*-GR24-regulated genes, AOS. This is perhaps unsurprising as the effect of SL on bud outgrowth is likely to differ between intact and decapitated plants due to the many changes that occur within a plant after decapitation (Dun et al., 2009), including increased sucrose levels (Mason et al., 2014), which has been shown to prevent SL inhibition of bud outgrowth in rose and pea (Bertheloot et al., 2020).

The SL transcriptome studies performed in intact plants, including ours, identified only a handful of differentially expressed genes after SL treatment compared to other hormones such as CK and ABA which have been shown to regulate hundreds, or even thousands of genes (Nemhauser et al., 2006; Bhargava et al., 2013). This implies that either these small number of genes are sufficient to control the phenotypes that SL regulates, or that SL responses may also be regulated by other processes within the cell, for example post-transcriptional, translational, or post-translational regulation, such as regulation of PIN1 polarisation (Shinohara et al., 2013). In support of this, we identified one gene co-expression module (Module 4) significantly associated with *rac*-GR24 treatment that was enriched in the GO terms translation, RNA methylation and ribosome biogenesis, and another (Module 6) enriched in rRNA processing and regulation of translation (Supplementary Dataset 3). In addition, we identified one SL-regulated gene, PsSKP2A, whose homolog in Arabidopsis is involved in post-translational protein regulation (del Pozo et al., 2002; del Pozo et al., 2006).

Another explanation for the relative paucity of DE genes identified in our study may be explained by the variability within the tissue samples collected as evidenced by the high degree of variation in gene expression we observed. Luo et al. (2019) proposed that SL repression of bud outgrowth occurs at a specific stage of bud development in rice, the P6 stage, as this was the stage at which WT and SL mutant buds began to differ phenotypically. Although we harvested buds from plants of a uniform age, the developmental stage of the buds may have differed between plants, and so the transcriptional responses of the buds to SL may also have differed.

### Gene co-expression analysis identifies a number of key co-expression modules regulated by SL in pea buds

We identified nine co-expression networks that were significantly correlated with *rac*-GR24 treatment. GO term enrichment analysis of the genes present in each of the nine modules identified key biological processes associated with *rac*-GR24 treatment. Not unexpectedly the enriched GO terms we identified after *rac*-GR24 treatment had a high degree of overlap with those identified as associated with increased bud dormancy in WT rice buds and with reduced expression in *tb1* buds (Dong et al., 2019).

Of particular note was the number of hormone biosynthesis and signalling GO terms enriched in many of the modules (Supplementary Dataset 3) suggesting that SL signalling may interact with other hormone pathways. Consistent with our results, some of these hormonal interactions have previously been identified; for example, ABA and JA biosynthesis and signalling has previously been implicated in SL (Luo et al., 2019) and BRC1 (González-Grandío et al., 2013; González-Grandío et al., 2017) regulation of bud outgrowth. We have further identified enrichment of GO terms associated with salicylic acid (SA) signalling and catabolic process, and ethylene biosynthesis and signalling in Modules 15 and 19 (Supplementary Dataset 3). There is some evidence that both ethylene (Tayama and Carver, 1990; Hayashi et al., 2001; Tsuchisaka et al., 2009) and SA (El-Esawi et al., 2017) positively regulate shoot branching. Future work should, therefore, examine further whether SL regulation of shoot branching involves ethylene or SA pathways. Interestingly, we did not observe any transcriptional changes in response to *rac*-GR24 in pea homologs of genes related to auxin transport as previously reported in rice (Sun et al., 2014; Xu et al., 2015). This suggests that SL-mediated regulation of auxin transport, at least initially, does not occur through transcriptional regulation, and instead must occur through post-transcriptional or translation regulation such as regulation of localisation of the PIN1 protein as previously demonstrated by Shinohara et al. (2013). SL has also been previously shown to regulate CK biosynthesis (Chen et al., 2013), metabolism (Duan et al., 2019) and signalling (Zha et al., 2019) genes. We did not identify any enrichment of CK genes in our WGCNA modules, but we did identify two CK-related genes as putative SL-responsive genes; comp84614_c0 homologous to CKX1 and comp73908_c1 homologous to ARR2, which were both positively regulated by SL (Supplementary Dataset 1).

Sugars are another key signal regulating shoot branching in plants (Mason et al., 2014; Barbier et al., 2019b). We identified two modules (6 and 9) in the WGCNA analysis with significant enrichment of GO terms associated with sugars (Figure 4D and Supplementary Dataset 3). In addition, two of the 15 SL-regulated genes, *PsCRCT* and *PsPSPE*, have putative roles in sucrose metabolism and transport. PsPSPE is a putative sugar phosphate exchanger based on homology to AT4G32480 (Table 1), while the closest homolog of comp71047_c1 in rice is a gene called *CRCT* (Os05g51690/Os05g0595300). In response to external factors such as CO_2_ and sucrose, CRCT was found to decrease the amount of available sucrose and increase starch content, likely by regulating starch synthesis genes such as *AGPL, AGPS* and *PhoI* (Morita et al., 2015). Interestingly, overexpressing *CRCT* in rice led to a slight decrease in the number of tillers (Morita et al., 2015) which suggests a putative role for CRCT in the regulation of shoot branching perhaps by controlling sucrose and starch content in buds.

### A high degree of interaction occurs between SL, CK and decapitation signalling in pea buds

Many studies have investigated the numerous interactions between the signalling pathways that regulate shoot branching. In this study we have provided further evidence for interactions between the SL signalling pathway and CK and decapitation by identifying novel genes regulated by all three signals in pea buds. As previously demonstrated, BRC1 is a key integrator of many of the signals that control axillary bud outgrowth (Barbier et al., 2019b), which we also demonstrate here with *PsBRC1* being upregulated by SL (Figure 1) and downregulated by CK (Figure S5) and decapitation (Figure S6) in pea buds. We further show that other SL responsive genes are also regulated by CK and decapitation mostly in an antagonistic manner to SL (Figure 3) as seen for *PsBRC1*. In fact, all of the BRC1-dependent genes that were upregulated by SL were also downregulated by CK and decapitation, while conversely most of the BRC1-independent genes were not antagonistically regulated by SL, CK and decapitation (Figure 6). This would suggest that SL, CK and decapitation converge on or before BRC1 to regulate downstream targets.

Interestingly, we also found that CK and decapitation upregulate expression of some *PsSMXLs* that likely act as upstream suppressors of BRC1 in the SL signalling pathway. This suggests that PsSMXLs are not specific to the SL signalling pathway, and instead may act as integrators of multiple signals that regulate shoot branching, similar to BRC1. While SL upregulation of *PsSMXL7* and *PsSMXL8* is likely due to feedback upregulation after protein degradation (Jiang et al., 2013; Song et al., 2017), upregulation of these genes by CK and/or decapitation is more likely to be a direct upregulation of these genes to increase protein levels. If this is indeed the case, then CK and decapitation regulation of *PsBRC1* may occur through upregulation of *PsSMXLs* transcript and protein levels. However, Dun et al. (2012) demonstrated that CK regulation of *PsBRC1* could occur in the absence of new protein synthesis (using cyclohexamide (CHX) treatment), which would indicate that CK regulation of PsBRC1 also occurs directly. Therefore, we suggest that decapitation and CK regulation of *PsBRC1* likely occurs both directly and indirectly through upregulation of *PsSMXL* expression (Figure 6).

## METHODS

### Plant material, growth conditions and measurements

The *Pisum sativum* plants used in this study include WT (L107), *rms1-2 (rms1-2T), rms4-1* (K164), *rms3-2* (K564) and *rms5-3* (BL298) genotypes previously obtained in cv Torsdag (Dun et al., 2012; de Saint Germain et al., 2016), as well as *rms1-10* (M3T-884) and *rms1-l0 PsBRC1^Té^* genotypes previously obtained in cv Térèse (Braun et al., 2012). Seeds were planted 2 per 2L pot in B2 potting mix (Green Fingers, Nerang, Australia; Figure S1A and S6), 4 per 2L pot in UQ23 potting mix (70% composted pine bark 0-5mm, 30% cocoa peat, mineral fertiliser; Figures 2, S2 and S5), and 4 (Figure 1) or 5 (Figure S1B) per 2L pot in EcoZ Plus potting mix (Green Fingers, Nerang, Australia). Seedlings were grown under 18h photoperiod until 8-days-old, unless otherwise specified. For Figure 5, pea plants were grown individually in glasshouse (23°C day/ 15°C night) under a 16-h photoperiod (the natural daylength was extended or supplemented during the day when necessary using sodium lamps) in 0.4 L pots filled with clay pellets, peat, and soil (1:1:1) supplied regularly with nutrient solution. Nodes were counted starting from the cotyledonary node as node 0. Bud length was measured with digital callipers (Figure S2) and with ImageJ using digital photographs taken of the bud every 30 minutes as described by Mason et al. (2014; Figure 1 A).

*Arabidopsis thaliana* Col-0, *d27-1* and *max4-1* were grown as described in Brewer et al. (2015). T-DNA Arabidopsis mutants generated by the Salk Institute Genomic Analysis Laboratory (Alonso et al., 2003) were obtained from NASC. T-DNA homozygous lines were verified by PCR on DNA extracted from leaf tissue using the primers outlined in Table S3. The number of rosette leaves was counted just after bolting of the main shoot, and the number of rosette branches longer than 5 mm was counted 28 or 45 days after bolting.

### Hormone and other treatments in pea

Unless otherwise specified, pea node 2 buds were treated with 10 μL of a water solution containing 0.01% Tween-20, 1% PEG 1450, 6.25% EtOH and differing concentrations of hormones as described, with the exceptions of Figure S1B which also had 37.5% EtOH and 0.1% DMSO in solution and Figure 5 which contained 2% PEG 1450, 50% EtOH and 0.4% DMSO. For *rac*-GR24 and GR24^5DS^ treatments 1 μM in acetone (0.1% final concentration) was used, and for BA (benzyladenine) treatments 100 μM in DMSO (0.1% final concentration) was used. For decapitation treatments (Figure S6) the shoot tip was removed at internode 5.

### Identification of differentially expressed transcripts

The pea bud transcriptome assembled by Kerr et al. (2017) was used as a reference, and the reads sequenced in Kerr et al. (2017) were used for differential expression analysis to identify transcripts that were differentially expressed in *rac*-GR24-treated pea buds compared to mock-treated pea buds. The three replicates for each sample were harvested over a 1h 40m time window to ensure that any variation due to diurnal changes as identified in Kerr et al (2017) were equally represented in all samples.

RSEM (version 2013-04-12) (Li and Dewey, 2011) was used to calculate read counts for all samples using the default parameters, except that the reference file was produced using nopolyA.

edgeR (Bioconductor version 3.2.4) (Robinson et al., 2010) was run to determine the differentially expressed transcripts using only the read counts for read 1 of the samples. The transcripts were first filtered to remove any transcripts with less than ten read counts in total. Pairwise comparisons were then made between the control and *rac*-GR24 treated samples at each of the four time points (1, 2, 4 and 6 hrs following treatment). A generalised linear model (GLM) was used to compare the *rac*-GR24 treatment to the control treatment over all four time-points. To determine differentially expressed genes, an FDR threshold of 0.05 was used.

Differentially expressed genes were annotated with a pea gene from the recently assembled pea genome (Kreplak et al., 2019) by running BLASTN (Camacho et al., 2009) locally to align each gene in the pea transcriptome (from Kerr et al., 2017) to the CDS sequences from the pea genome (from Kreplak et al., 2019).

### WGCNA analysis

The variance-stabilized count data of the genes that had a total count of equal to or more than 10 across all treatment-time combinations were used for co-expression network analysis using WGCNA package 1.64.1 (Langfelder and Horvath, 2008). These selected genes were ranked by their variance across all samples and the 5000 most varying genes (10%) were selected for WGCNA analysis. A signed network with soft threshold power of 16 was created using WGCNA package version 1.64.1 (Langfelder and Horvath, 2008). During the process, outlier sample 25 was removed from analysis. Hub genes were identified based on intramodular connectivity. The network was visualized using Cytoscape version 3.5.1 (Shannon et al., 2003).

### Phylogenetic analyses

Transcript sequences for each DE gene was obtained from Kerr et al. (2017) and translated into predicted protein sequences using the ExPASy Translate tool (Gasteiger et al., 2003); if more than one sequence existed for each transcript then the sequence with the longest open reading frame (ORF) was used. Putative orthologs from *Arabidopsis thaliana, Medicago truncatula, Glycine max, Populus trichocarpa, Oryza sativa* and *Physcomitrella patens* were identified using OrthoDB (Kriventseva et al., 2015). Protein sequences were used to generate amino acid sequence similarity trees using the Maximum Likelihood method based on the Dayhoff matrix based model (Schwarz and Dayhoff, 1979), and were conducted in MEGA7 (Kumar et al., 2016).

### Gene expression analyses

For qRT-PCR in Figures 1, 1B, S5 and S6, total RNA was isolated and quantified as described by Braun et al. (2012). For Figures 2 and S2, total RNA was isolated using a CTAB protocol (Barbier et al., 2019a). RNA was quantified on a NanoVue Plus and migrated on gels to determine RNA degradation. For Figure 5, total RNA was isolated using the RNeasy plant mini kit (Qiagen) following manufacturer’s instructions. DNase treatment was performed to remove DNA using the RNase-Free DNase Set (Qiagen) eluted in 50 mL of RNase-free water. RNA was quantified using NanoDrop 1000.

For all qRT-PCR, except figure 5, cDNA was synthesised from RNA using the iScript™ reverse transcription supermix (BioRad) as per the manufacturer’s instructions. The cDNA was diluted to 0.5 ng/μL for qRT-PCR. For Figure 5, cDNA was synthesized from 2 μg of total RNA using 50 units of RevertAid H Moloney murine leukemia virus reverse transcriptase in 30 μL following the manufacturer’s instructions with poly(T)18 primer. cDNA was diluted 10 times before subsequent analysis.

qRT-PCR was performed and analysed as previously described (Mason et al., 2014). *Ef1*α, *GADPH* and *TUB2* were used as reference genes. Primer sequences were designed using Primer3 software (Untergasser et al., 2012) based on transcript sequences from Kerr et al. (2017) and can be found in Table S4. qRT-PCR products were sequenced to confirm primer amplification of the correct product.

The heat map was constructed with Treeview (Saldanha, 2004) using log fold changes in gene expression of qRT-PCR data with clustering performed by Cluster3 (de Hoon et al., 2004).

### Statistical analyses

Statistical analyses were performed in R (R Core Team, 2013) using the car package (Fox and Weisberg, 2019). Levene’s test and the Shapiro-Wilk test were used to determine if data fit the homogeneity of variance and normal distribution assumptions, respectively. If necessary extreme outliers were removed from the datasets. Depending on the dataset, either a Student t-test, oneway ANOVA or two-way ANOVA was performed, with Tukey’s post-hoc test if appropriate. If the homogeneity of variance assumption was violated then Welch’s test was performed.

## SUPPLEMENTARY DATA FILES

Figure S1: SL inhibits bud growth from at least 6.5 hrs onwards and regulates gene expression within 1 hr

Figure S2: GR24^5DS^, but not KAR_1_, can significantly inhibit node 2 bud growth

Figure S3: Characterisation of branching phenotype in mutants of SL-regulated genes

Figure S4: Module-trait relationships identified in the top 5000 variable genes in pea buds

Figure S5: CK regulates many SL-regulated genes

Figure S6: Decapitation regulates many SL-regulated genes

Table S1: SL-regulated genes at different FDR thresholds

Table S2: Summary of shared genes in pea and Arabidopsis networks

Table S3: List of primer sequences used to verify Arabidopsis TDNA homozygous lines

Table S4: List of primer sequences used for qRT-PCR

Supplementary Dataset 1: List of potential strigolactone-regulated genes identified using RNA-Seq at 1, 2, 4 and 6 hrs following *rac*-GR24 treatment

Supplementary Dataset 2: SKP2A may negatively regulate branching in Arabidopsis

Supplementary Dataset 3: Biological processes GO term enrichment in nine modules significantly correlated with *rac*-GR24 treatment and/or length of *rac*-GR24 treatment

Supplementary Dataset 4: Top five hub genes for each of the nine modules significantly correlated with *rac*-GR24 treatment and/or length of *rac*-GR24 treatment

## ACKNOWLEDGEMENTS

We would like to acknowledge Associate Professor Jessica Mar for help with the RNA-Seq GLM analysis and Associate Professor Christopher McErlean for supplying the GR24^5DS^. The research was supported by funding from The University of Queensland and the Australian Research Council.

## AUTHOR CONTRIBUTIONS

SCK, MT and CB designed the research. SCK and AdSG performed the research. ID performed the WGCNA. MM and ED provided cDNA. SCK and MT analysed the data. SCK and MT wrote the manuscript. All co-authors read and commented on the manuscript.

