## Supplementary Figures and Tables for "Hormonal regulation of the BRC1-dependent strigolactone transcriptome involved in shoot branching responses"

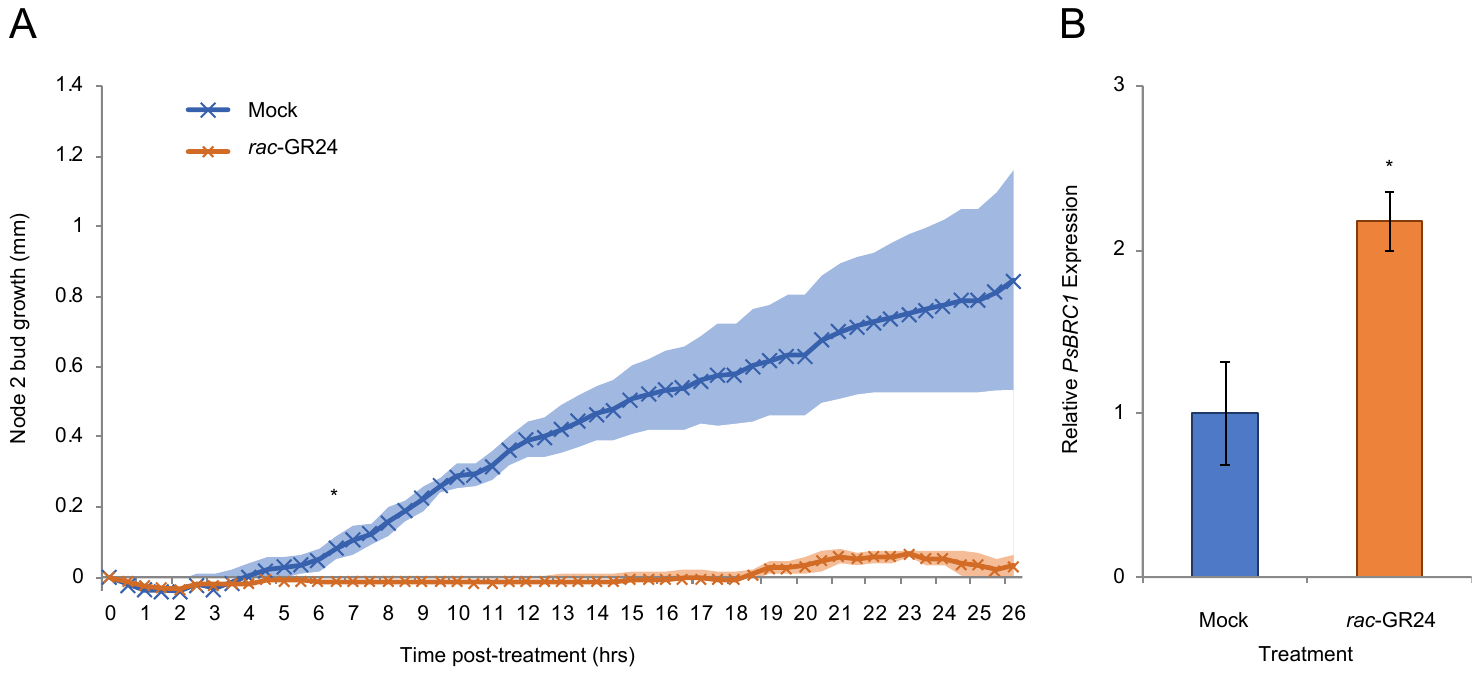

**Figure S1** **SL inhibits bud growth from at least 6.5 hrs onwards and regulates gene expression within 1 hr. A)** The bud at node 2 of 8-day-old *rms5-3 Pisum sativum* plants was treated with a solution containing 0 (Blue) or 1 µM (Orange) of the synthetic strigolactone, *rac*-GR24. Bud growth was captured using time-lapse photography every 30 mins and measured using ImageJ. Data are means ± SE (n = 4). * denotes the time at which the mock and *rac-*GR24 treatments become significantly different, namely 6.5 hrs (Student t-test; P<0.05). **B)** The bud at node 2 of 7-day-old *rms1-2 Pisum sativum* plants was treated for 1 hr with a solution containing 0 (Blue) or 1 µM (Orange) *rac*-GR24. Expression of *PsBRC1* is represented relative to the mock treatment; *EF1α* was used as an internal reference gene. Data are means ± SE (n = 3 pools of ~28 buds). * denotes means significantly different from the mock treatment (Student t-test; P<0.05).

|  |  | **FDR<0.05** | | | | **FDR<0.5** | | | |
| --- | --- | --- | --- | --- | --- | --- | --- | --- | --- |
| **Time point** | **Analysis** | **Down-regulated transcripts** | **Up-regulated transcripts** | **Total DE transcripts** | **Transcripts validated by qRT-PCR** | **Down-regulated transcripts** | **Up-regulated transcripts** | **Total DE transcripts** | **Transcripts validated by qRT-PCR** |
| 1hr | Pairwise | 0 | 0 | 0 | n/a | 0 | 0 | 0 | n/a |
| 2hr | Pairwise | 0 | 0 | 0 | n/a | 1 | 0 | 1 | 0 (0%) |
| 4hr | Pairwise | 2 | 3 | 5 | 3 (60%) | 3 | 9 | 12 | 7 (58%) |
| 6hr | Pairwise | 0 | 5 | 5 | 5 (100%) | 5 | 22 | 27 | 17 (63%) |
| Over-time | GLM | 1 | 9 | 10 | 9 (90%) | 104 | 182 | 286 | 16 (55%) |
| **Total unique DE transcripts** | | **3** | **11** | **14** | **11 (79%)** | **112** | **194** | **306** | **26 (52%)** |

**Table S1** **SL-regulated genes at different FDR thresholds**. Summary of the number of transcripts identified as differentially expressed in pea buds treated with 1 µM rac-GR24 using a FDR threshold of <0.05 and FDR<0.5.

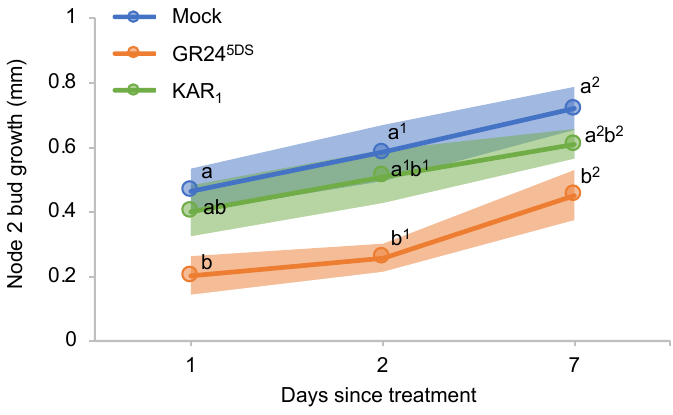

**Figure S2** **GR24^5DS^, but not KAR_1_, can significantly inhibit node 2 bud growth**. The bud at node 2 of 9-day-old *rms1-2* *Pisum sativum* plants was treated with a solution containing 0 (Blue) or 1 µM (Orange) GR24^5DS^ or 1 µM (Green) KAR_1_. Bud length (mm) was measured before and 1, 2 and 7 days following treatment to calculate bud growth. Data are mean ± SE (n = 14-15). Data at each time point were separately analysed using a one-way ANOVA with Tukey comparisons of means; different letters represent statistical differences of P<0.05.

A

B

|  | Primary rosette branches |
| --- | --- |
| Col-0 | 5.83333 |
| max4 | 16.7273 |
| d27 | 11.7333 |
| aos | 4.91667 |
| dall2 | 5.63636 |
| nsh1 | 5.84615 |
| erf061 | 5.78571 |
| kuf1 | 5.8125 |
| skp2a | 7.84615 |
| tbl21 | 5.92308 |

WT^Col-0^	*srf1-1	srf 1-2*

|  | Primary rosette branches |
| --- | --- |
| Col-0 | 8.3125 |
| srf1-1 | 7.64286 |
| srf1-2 | 7.64286 |

**Figure S3** **Characterisation of branching phenotype in mutants of SL-regulated genes.** The number of rosette branches scored A) 45 days or B) 28 after bolting of the main stem of *Arabidopsis thaliana* mutant lines. Data are mean ± SE (n = 8-16). * denotes means significantly different from Col-0 (Student t-test; P<0.05). Data were analysed using a one-way ANOVA with Tukey comparisons of means, or Welch’s one-way test if homogeneity of variance assumption was violated; different letters represent statistical differences of P<0.05.

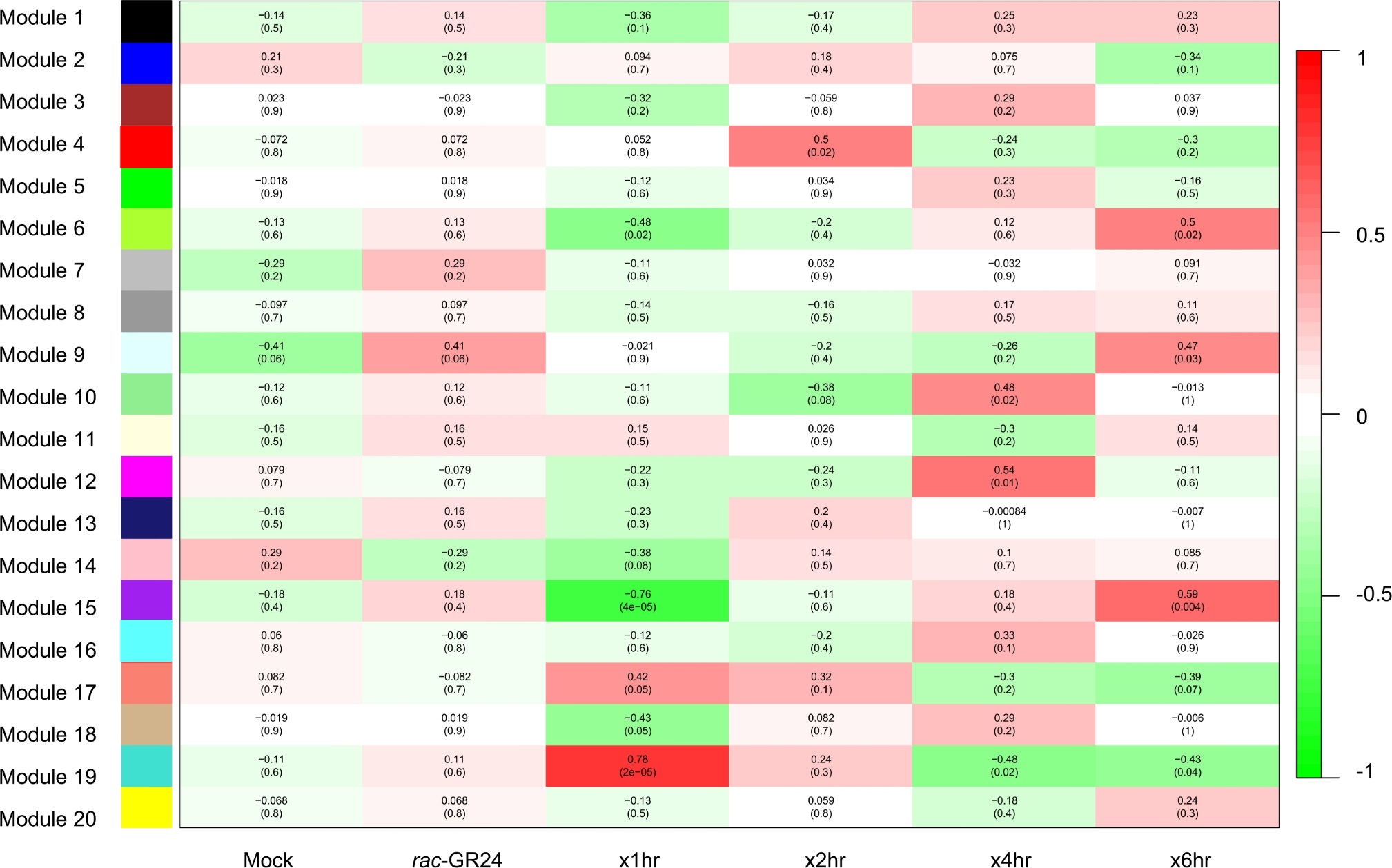

**Figure S4 Module-trait relationships identified in the top 5000 variable genes in pea buds.** Module-trait relationships of the 20 co-expression modules identified in the top 5000 most variables genes in pea buds comparing mock and *rac*-GR24 treatment over four treatment lengths.

**Table S2** **Summary of shared genes in pea and Arabidopsis networks.** Shared genes co-expressed with 13 *rac*-GR24 regulated genes in pea and Arabidopsis GeneMANIA networks (Warde-Farley et al., 2010).

| **Gene of interest** | | | **# genes co-expressed with gene of interest in pea network** | **# pea genes with AGI** | **# genes connected to gene of interest in Arabidopsis GeneMANIA network** | **# genes shared between pea and Arabidopsis networks** | **% shared genes present in pea network with AGI** | **% shared genes present in Arabidopsis network** | **Hyper-geometric P value** |
| --- | --- | --- | --- | --- | --- | --- | --- | --- | --- |
| **Pea comp** | **Gene name** | **AGI** |  |  |  |  |  |  |  |
| comp78442_c0 | PsBRC1 | AT3G18550 | 22 | 20 | 0 | 0 | 0% | n/a | 1 |
| comp114246_c0 | PsERF061 | AT1G64380 | 12 | 11 | 74 | 1 | 9% | 1% | 2.87E-02 |
| comp102807_c0 | PsAOS | AT5G42650 | 268 | 197 | 123 | 19 | 10% | 15% | 3.23E-20 |
| comp55781_c0 | PsPSPE | AT2G20670 | 94 | 70 | 85 | 9 | 13% | 11% | 8.69E-13 |
| comp70495_c0 | PsNCED2 | AT3G14440 | 81 | 64 | 61 | 2 | 3% | 3% | 8.45E-03 |
| comp70728_c0 | PsNSH1 | AT2G36310 | 64 | 55 | 93 | 6 | 11% | 6% | 3.05E-08 |
| comp81043_c1 | PsSRF2 | AT5G06820 | 65 | 52 | 0 | 0 | 0% | n/a | 1 |
| comp87415_c1 | PsSKP2A | AT1G21410 | 32 | 27 | 68 | 6 | 22% | 9% | 4.98E-11 |
| comp86930_c0 | PsSMXL8 | AT2G40130 | 21 | 20 | 22 | 1 | 5% | 5% | 1.57E-02 |
| comp56263_c0 | PsDALL2 | AT1G51440 | 12 | 11 | 39 | 2 | 18% | 5% | 1.05E-04 |
| comp71047_c1 | PsCRCT | AT5G53420 | 18 | 15 | 51 | 0 | 0% | 0% | 9.73E-01 |
| comp93400_c0 | PsMIOX4 | AT4G26260 | 25 | 22 | 30 | 0 | 0% | 0% | 9.76E-01 |
| comp54718_c1 | PsRS6 | AT5G20250 | 60 | 46 | 80 | 5 | 11% | 6% | 2.19E-07 |
| **All 13 genes** | | | **434** | **320** | **434** | **36** | **11%** | **8%** | **2.16E-20** |

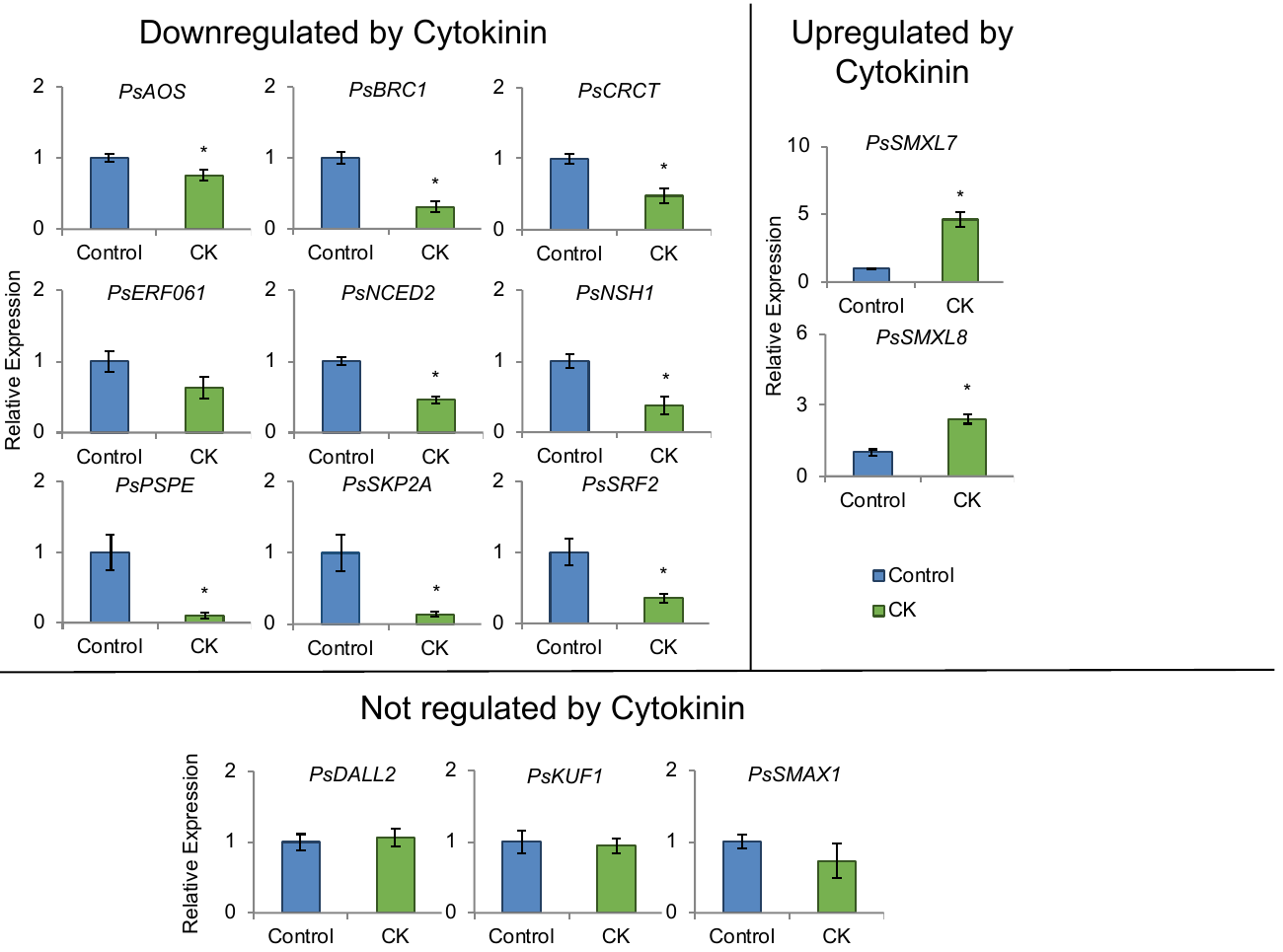

**Figure S5** **CK regulates many SL-regulated genes.** The bud at node 2 of 8-day-old Torsdag wild type (L107) *Pisum sativum* seedlings was treated for 2 hrs with a solution containing 0 (Blue) or 100 µM (Green; CK) benzyladenine (BA). Expression is represented relative to the mock treatment and was normalized against the geomean of three internal reference genes: *EF1α*, *GADPH* and *TUB2*. Data are means ± SE (n = 3 pools of ~60 plants). * denotes means significantly different from the mock treatment (Student t-test; P<0.05).

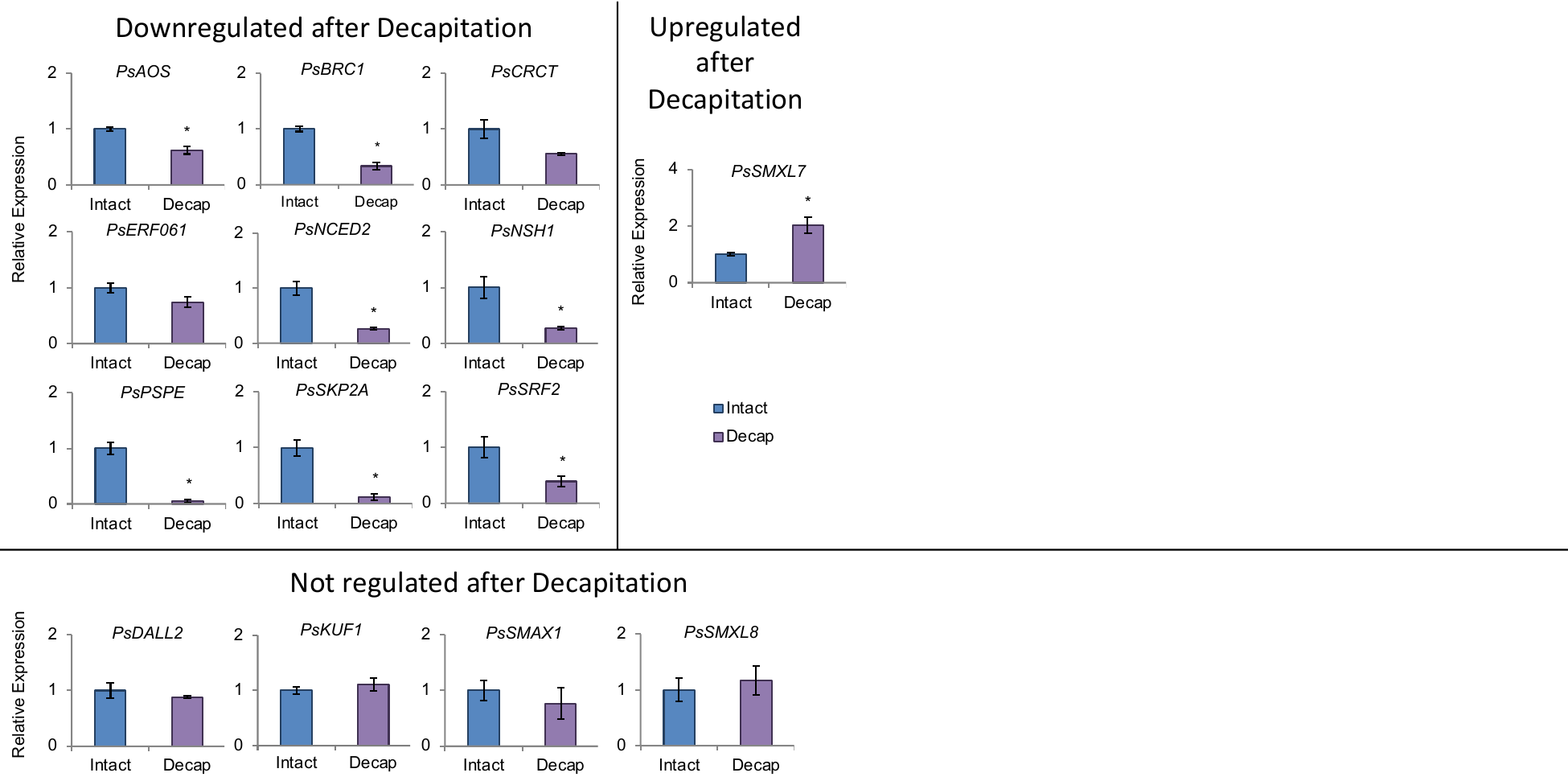

**Figure S6** **Decapitation regulates many SL-regulated genes.** Torsdag wild type (L107) *Pisum sativum* seedlings with 5 leaves expanded were left intact (Blue) or decapitated (Purple; Decap) at internode 5, and node 2 bud tissue was harvested 2 hrs later. Expression is represented as relative to Intact and was normalized against the geomean of three internal reference genes: *EF1α*, *GADPH* and *TUB2*. Data are means ± SE (n = 3 pools of ~20 plants). * denotes means significantly different from the Intact (Student t-test; P<0.05).

**Table S3** List of primer sequences used to verify Arabidopsis TDNA homozygous lines

| SALK line | Gene | Forward sequence (5’ – 3’) | Reverse sequence (5’ – 3’) | TDNA primer | TDNA sequence (5’ – 3’) |
| --- | --- | --- | --- | --- | --- |
| SALK_017756 (aos-1) | AT5G42650 (AOS) | CGACGAGAAATTAACGGAGC | GGAACTAACCGGAGGCTACC | LBb1.3 | ATTTTGCCGATTTCGGAAC |
| SALK_012432 (dall2-1) | AT1G51440 (DALL2) | TTATTACCCATCCACGATCCC | ATGACTATGTCACGTCTCCCG | LBb1.3 | ATTTTGCCGATTTCGGAAC |
| SALK_087511 (erf061-1) | AT1G64380 (ERF061) | AGTGAGGCAGAGACATTGGG | TCTCTTCTTACCTTCCCCACC | LBb1.3 | ATTTTGCCGATTTCGGAAC |
| SALK_206003 (kuf1-1) | AT1G31350 (KUF1) | CTTGCGATGTAGATAGCTCCG | TCTGTTCACCGGTTCTTTTTC | LBb1.3 | ATTTTGCCGATTTCGGAAC |
| SM_3_39680 (nsh1-1) | AT2G36310 (NSH1) | AAATCAGGCCGTACTACAGCC | AACTTTTGGAATGCGTCTGTG | Spm32 | TACGAATAAGAGCGTCCATTTTAGAGTGA |
| GABI-293D12 (skp2a-1) | AT1G21410 (SKP2A) | TGCAGACTTTAAATCTAAGGCAGG | ATGTTTCTGCAATAGTATAGCCCC | o3144/35St (pAC161) | GTGGATTGATGTGATATCTCC |
| GT_5_7628 (skp2a-2) | AT1G21410 (SKP2A) | GCCTACCTAGCCAAATGACATC | AGGATCCTCATCAGAAGCTCC | Ds3-1 | ACCCGACCGGATCGTATCGGT |
| SALK_034496 (tbl21-1) | AT5G15890 (TBL21) | GAGCTTCTTCGTTTGGTACCC | GAGCTTCTTCGTTTGGTACCC | LBb1.3 | ATTTTGCCGATTTCGGAAC |
| SALK_204435  (srf1-1) | AT5G06820 (SRF2) | TGGCAGAATTCGAGAATGAAC | TGAGTAGCGTTAGGAGGCAAG | LBb1.3 | ATTTTGCCGATTTCGGAAC |
| SAIL_407_E02  (srf1-2) | AT5G06820 (SRF2) | TAGTATCCTCACCCAGGCATG | GTGAGCAGAATTGTAGCGAGG | LB1 | GCCTTTTCAGAAATGGATAAATAGCCTTGCTTCC |

**Table S4** List of primer sequences used for qRT-PCR

| Gene | Forward sequence (5’ – 3’) | Reverse sequence (5’ – 3’) |
| --- | --- | --- |
| PsEF1α | TGTGCCAGTGGGACGTGTTG | CTCGTGGTGCATCTCAACGG |
| Ps-GADPH | TCGGACTTCAGGGATGTGTATT | GCTGGTGCGGAGTTTATCTG |
| Ps-TUB2 | AGATGGCTTCAACTTTCATTGG | GCTCTCGGCTTCGGTGA |
| comp101556_c0 | GAGCAACATGGTGTCACTGG | TCATATCCCTCAGGTCCTTCC |
| comp102807_c0 (PsAOS) | CGGTACAGCAGGTTCAATCC | TTTCCGACGAAATCAGAACC |
| comp114246_c0 (PsERF061) | CATCAAACACCACCAACACC | CGAAGATAAACCGAGGAACC |
| comp209307_c0 | AATTGGAGACTCGCAAAAGG | AATGGACTTCATGCCTGTGG |
| comp28144_c1 | CGCACGTCCACTATTTTTCC | TGTGTATCCCAAGCTGTAGGC |
| comp28866_c0 | GTACTGCACATTCCGATTGC | CACTCCACCATGAGTTTTGC |
| comp32609_c0 | ATAGGCTCGATGGTGTGTCC | CTACACGGACCGTCAGATCC |
| comp36904_c0 | CCCCTTGAATTTCCAGTTACC | GGGGACAAAGAAAGAGAAGG |
| comp378909_c0 (PsLDP) | TCTCTGATTCGGCTTCAAGG | CAATCCATTCCACCTTTTGC |
| comp511744_c0 | TTCAGCCATAAACCCTGACC | CCTCGCTTACCTTCTCATTGC |
| comp54718_c1 (PsRS6) | CGTGCCTGCTTTCTTCTTTC | AAAGCAGCGGTTAGGGTTTC |
| comp55397_c0 (PsFH18) | CCAAACTTTGATGGGAAAGG | CCCTCAATCACGACATCACC |
| comp55543_c0 | GGCTTGGTTGTTGTTGTTAGG | CAAGCATTTTGGTTCAATGC |
| comp55781_c0 (PsPSPE) | TCAAGGTGCCTTCAAAATGC | TGGTTTCTGTTTGCTTGTGC |
| comp56263_c0 (PsDALL2) | GATGGGATACATCGCTGTCG | TGGGTCATTTCTGAAGTTTGC |
| comp62406_c0 | CTATGGAGGAGGGTTGTTGG | CTGGCTGTGGTATCATGTGC |
| comp65268_c0 | GACCAACACTACCCTGAAATTAAC | CATAGCGAGGGTGTACATTGG |
| comp68995_c1 | AGAAGCGCAGACTTTGAAGG | AATCCTGATGGATCGTACCG |
| comp69582_c0 | GTTCCCTAGCGCATTTAAGC | CATTCCGCACTATCTTCTTGG |
| comp70464_c1 | CTTGGTGTTCGGTCTTTGGT | GAGCGCAATTCTCTTCCATC |
| comp70478_c0 | CAAACTCCAAAGGCCAATGT | TGTAAACGAGGATCAACGTCA |
| comp70495_c0 (PsNCED2) | AGGCATTGACGGTGTTTACC | AGCGGCAAGAGTAACTGACG |
| comp70728_c0 (PsNSH1) | CCAAAAACCGTTGTCAAACC | TGATACTGACCCTGGGATCG |
| comp71047_c1 (PsCMP) | TGGGATCCAGATCAATGTCC | AAAACCCAGAAGCTCAATGC |
| comp71172_c0 | CGGCAAAGAGGATATTCACG | TCACACAAGTCGGTTGAAGG |
| comp72889_c0 | ATTGCTTGTGTTGCTTGTGC | GTTGGAGGGATCAAGGAAGC |
| comp73484_c0 (PsTBL21) | TTATTCCATTCGCCGTTCTC | TACCGAACCGCTCTCAAAAC |
| comp74214_c2 (PsOSB1) | ACACTTGCCAAAGGTGAAGC | TTGTTTGGCTCCTTACATTGG |
| comp74856_c2 (PsGH3.3) | CTCCATTCCCAAACAACACC | CGAGTGTGGTGGAGTACACG |
| comp78374_c1 (PsSMXL7) | ATCAAAAGGTTGCGAAAACG | AAGGTTCCAAGCTCCATTCC |
| comp78442_c0 (PsBRC1) | TCTGCAGGTACACAAACTGTGA | TCATGTTCCTTTGCTTTTTGA |
| comp78560_c0 | GCGAAAGATGGTGAACAACC | TGATCATCGTCAACAACTTGG |
| comp78689_c0 (PsSMAX1) | TCAGTGCAAATTAGGCAACG | TTGGCTTTGACCCTGATACC |
| comp79583_c0 | TGCTCTTATCCCCATAAATTGC | TGAAAAGTTGTGTTGCAGTGG |
| comp80029_c0 | TCGCTTTTCCACTCCTAACC | GTGTTTTCCTTTGGGGAACC |
| comp80781_c0 (PsKUF1) | GGATGAGATTGTCGGAGACG | ATCAAACACCTGGCAGATCC |
| comp81043_c1 (PsSRF2) | CAGAACCACAAAAGCAATGG | CTCTGTAAACAGGGCCAAGG |
| comp81735_c1 (PsDDP) | TTAAAGCCAGGGTTTTCAGG | TGAGCAAGAGGTGGAAAAGC |
| comp86446_c0 | GGGGAGTAACTTTGCGACAC | TTGCGAATATCGTCAGCATC |
| comp86903_c0 | TGGTGTTGTGCTTCTTGAGC | CTTTCCCGTTCTAACCAACG |
| comp86930_c0 (PsSMXL8) | TAATGTTTCTCCCGATCACG | CGAAAACCCGACTAACAACC |
| comp87019_c0 (PsCYS5) | GAGGTTTCGGGAGTGAAGC | CTCAGTACGACAAGCGAAGC |
| comp87415_c1 (PsSKP2A) | TTCATTGGCACAGAGCAAAG | AGCCTGAACAGCTGAAGGAG |
| comp88452_c2 | ATCAACCACCTCACTCTCGC | GGTCAAAAGTGGTTCCATGTCG |
| comp90222_c0 (PsALKR5) | CAGCATCGGATTGATACTCG | TCTGATTGTTGCAGCTGAGG |
| comp91910_c0 | TTCGTGAGAACCTCTTTTGG | GCTTTCGCGTATTCAGAACC |
| comp93400_c0 (PsMIOX4) | CCTCCCTCATTTTCTTCACG | CTCACTCCGGAAATCAATGC |
| AtActin | AGTGGTCGTACAACCGGTATTGT | GATGGCATGAGGAAGAGAGAAAC |
|  |  | GAGGAAGAGCATACCCCTCGTA |
|  |  | GAGGATAGCATGTGGAAGTGAGAA |
| AtSKP2A | TTTGAATCTTTGCGGATGTG | CAGAGGTCAAGGGTCCTGAG |
