## Supplementary Dataset 2 for "Hormonal regulation of the BRC1-dependent strigolactone transcriptome involved in shoot branching responses"

**Supplementary Dataset 2** SKP2A may negatively regulate branching in Arabidopsis

*Atskp2a-1* was the only mutant that displayed any significant alteration in branching phenotype in our assay, with two more rosette branches than the WT (Figure S7). Interestingly, we found that while *Atskp2a-1* had a slight increase in branching, *Atskp2a-2* had a slight decrease in branching (Supplementary Dataset 2 Figure 1C, D). Although these branching differences are only small, they were replicated in a separate experiment (Supplementary Dataset 2 Figure 1E). The two *Atskp2a* mutants have T-DNA insertions in different locations of the *AtSKP2A* gene; *Atskp2a-1* contains a T-DNA insertion in the third exon of *AtSKP2A*, while *Atskp2a-2* contains a T-DNA insertion in the 5’UTR of *AtSKP2A* (Supplementary Dataset 2 Figure 1A). To investigate whether the contrasting branching phenotype in the two *Atskp2a* mutants was due to differential expression of the *AtSKP2A* gene, we measured *AtSKP2A* gene expression in the two mutants using two different primer pairs. Indeed, the two *Atskp2a* mutant lines had different *AtSKP2A* expression patterns; *Atskp2a-1* had no significant change in expression of *AtSKP2A*, while *Atskp2a-2* had increased expression of *AtSKP2A* (Supplementary Dataset 2 Figure 1E). We hypothesise that the *Atskp2a-1* T-DNA insertion in the third exon of *AtSKP2A*, while not affecting *AtSKP2A* expression may affect AtSKP2A protein function. This is especially pertinent given that the T-DNA insertion is located in the leucine-rich repeat (LRR) domain (Supplementary Dataset 2 Figure 1B) which is likely important for protein-protein interactions (Bella et al., 2008). In fact, this domain has previously been shown to be important for SKP2A binding to auxin (Jurado et al., 2010). In contrast, we hypothesise that the *Atskp2a-2* T-DNA insertion in the 5’UTR of *AtSKP2A* while increasing *AtSKP2A* expression may not have any effect on AtSKP2A protein function. These hypotheses would indicate that AtSKP2A function is negatively correlated with branching and suggests a role for AtSKP2A in the regulation of shoot branching, but further investigations are needed to confirm these hypotheses.

**Bella J, Hindle KL, McEwan PA, Lovell SC** (2008) The leucine-rich repeat structure. Cell Mol Life Sci **65**: 2307–2333

**Jurado S, Abraham Z, Manzano C, López-Torrejón G, Pacios LF, Pozo JCD** (2010) The Arabidopsis Cell Cycle F-Box Protein SKP2A Binds to Auxin. Plant Cell **22**: 3891–3904

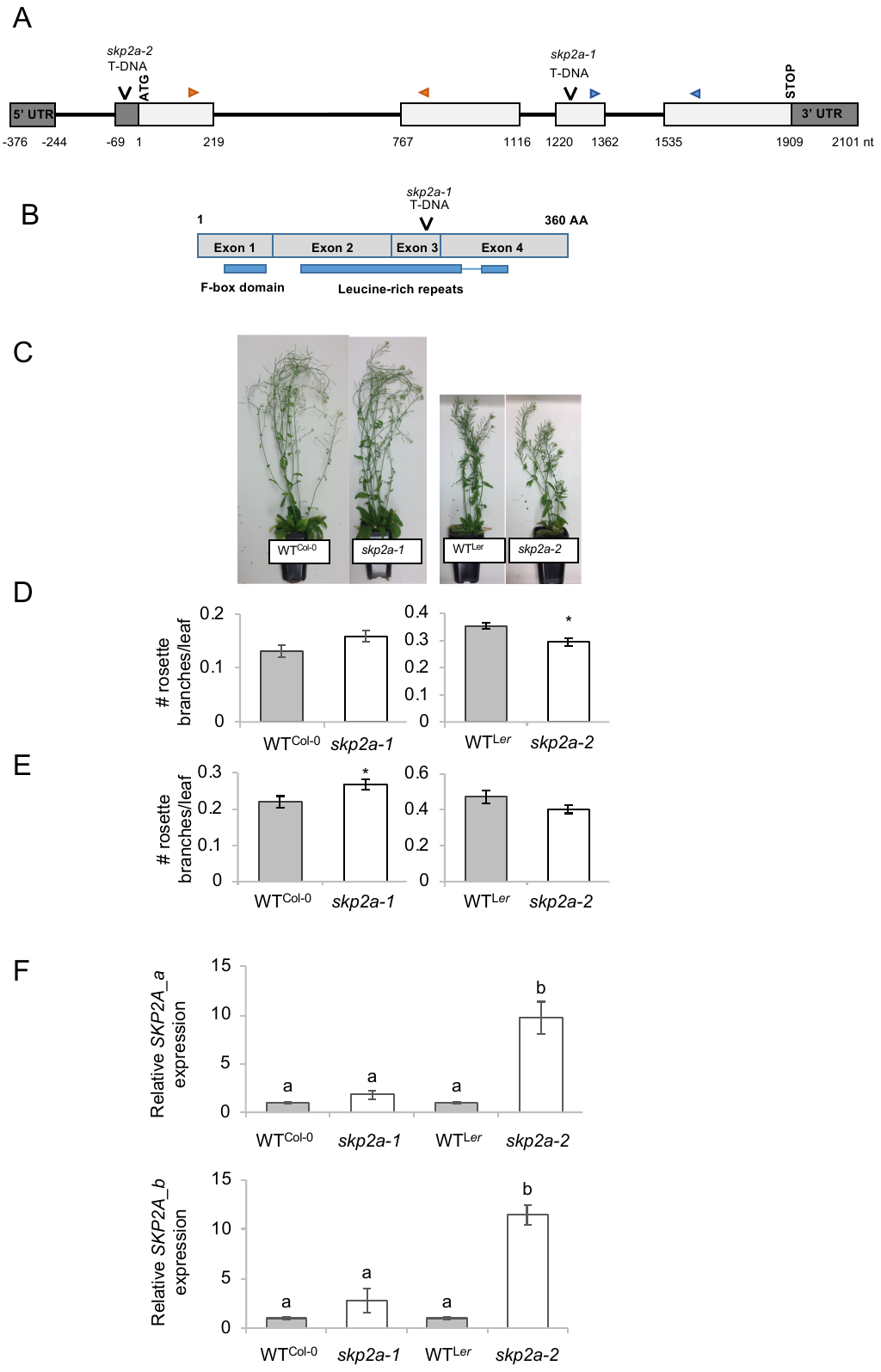

**Supplementary Dataset 2 Figure 1** **SKP2A may negatively regulate branching in Arabidopsis. A)** Structure of the *PsSKP2A* gene showing location of the two *skp2a* mutations. Bases are numbered from the start codon and are based on the genomic sequence; intron (thin line). The qRT-PCR primers locations are indicated by the coloured arrows (*SKP2A_a* orange ; *SKP2A_b* green). **B)** Structure of the PsSKP2A protein showing the location of the *skp2a-1* T-DNA insertion and the protein domains. **C-D)** The number of rosette branches > 5mm per rosette leaf was scored 28 days after the main stem had bolted on WT and *skp2a* mutant plants on a Columbia (Col-0) or Landsberg *erecta* (L*er*) background. Data are means ± SE (n = 18-20). * denotes means significantly different from the WT (Student t-test; P<0.05) **E)** The number of rosette branches > 5mm per rosette leaf was scored after plant senescence on WT and *skp2a* mutant plants on a Columbia (Col-0) or Landsberg *erecta* (L*er*) background. Data are means ± SE (n = 10-12). * denotes means significantly different from the WT (Student t-test; P<0.05). **F)** Expression of *SKP2A* in 3-week old *Arabidopsis thaliana* WT and *skp2a* seedlings on a Columbia (Col-0) or Landsberg *erecta* (L*er*) background. Expression is represented relative to each respective WT; Actin was used as an internal reference gene. Data are means ± SE (n = 3 pools of 5 seedlings). Data were analysed using a one-way ANOVA with Tukey comparisons of means, or Welch’s one-way test if homogeneity of variance assumption was violated; different letters represent statistical differences of P<0.05.
